# Bidirectional regulation between AP-1 and SUMO genes modulates inflammatory signalling during *Salmonella* Typhimurium infection

**DOI:** 10.1101/2022.03.18.484857

**Authors:** Pharvendra Kumar, Amarendranath Soory, Salman Ahmad Mustfa, Dipanka Tanu Sarmah, Samrat Chatterjee, Guillaume Bossis, Girish S Ratnaparkhi, C. V. Srikanth

**Affiliations:** Regional Centre for Biotechnology, 3rd milestone Gurgaon Faridabad Expressway, Faridabad, India; Kalinga Institute of Industrial Technology, Bhubaneshwar, India; Indian Institute of Science Education and Research, Pune, India; Kings College London, UK; Translational Health Science and Technology Institute, 3rd milestone Gurgaon Faridabad Expressway, Faridabad, India; Institut de Génétique Moléculaire de Montpellier (IGMM), Univ Montpellier, CNRS, Montpellier, France

## Abstract

Gram-negative bacterium *Salmonella* Typhimurium (*STm*) is the causative agent of gastroenteritis. Among the various gut pathogens, *STm* is still one of the most frequent culprits posing a significant health challenge. *<u>STm</u>* utilizes its effector proteins to highjack host cell processes. Alteration of SUMOylation, a post-translational modification mechanism, is one such change caused by *STm. STm* mediated simultaneous downregulation of SUMO-pathway genes, Ubc9 and PIAS1, is required for an efficient infection. In the present study, the regulation of SUMO pathway genes during *STm* infection was investigated. Promoters of both UBC9 and PIAS1, were seen to harbor binding motifs of AP-1, Activator protein-1 (c-Jun:c-Fos heterodimers or c-Jun:c-Jun homodimers). Using electrophoretic mobility shift assays, a direct binding of c-Fos to the identified motifs was observed. Perturbation of c-Fos led to changes in expression of Ubc9 and PIAS1, while its SUMO-modifications resulted in differential regulation of its target genes. In line with this, *STm* infection of fibroblasts with SUMOylation deficient c-Fos (c-FOS-KO^SUMO-def-FOS^) resulted in uncontrolled activation of target genes, as revealed by 3’mRNA-Seq analysis and mathematical modelling, resulting in massive activation of inflammatory pathways. Infection of c-FOS-KO^SUMO-def-FOS^ cells favored *STm* replication, indicating misdirected immune mechanisms in these cells. Finally, chromatin Immuno-precipitation assays confirmed a context dependent differential binding and release of AP-1 to/from target genes due to its Phosphorylation and SUMOylation respectively. Overall, our data point towards existence of a bidirectional cross-talk between c-Fos and the SUMO pathway and highlighting its importance in AP-1 function relevant to STm infections and beyond.

**Author summary:** Food borne infections caused *Salmonella* Typhimurium pose a major health challenge in developing and developed world. Unfortunately, many aspects of Salmonella-host crosstalk still remain unknown. In the current work, using sophisticated computational tools along with cell culture experiments and mathematical modeling, we demonstrate how *Salmonella* controls SUMOylation, a post-translational modification (PTM) pathway of host. SUMOylation governs fundamental processes of the host cell, and its alteration is required for a successful *Salmonella* infection. We show that SUMO-pathway genes, Ubc9 and Pias1, are direct target genes of AP-1 transcription factor. C-Fos, a component of AP-1 transcriptionally regulates SUMO-genes by binding to their promoters. During *Salmonella* infection, a selective activation of target genes of c-Fos was observed. The selective regulation of target genes relied on c-fos PTMs. Experimental perturbation of c-Fos PTMs led to global transcriptional dysregulation including immune hyperactivation. Thus, we show existence of a complex interplay between the SUMO-pathway genes and AP-1 transcription factors which mediate selective gene regulation during *Salmonella* infection.

## Introduction

The gram-negative pathogen *Salmonella enterica* serovar Typhimurium (*STm*) is a causative agent of gastroenteritis in humans. The disease occurs due to consumption of contaminated food or water, leading to a localized infection in the intestine, which triggers acute diarrhea, accompanied by abdominal cramps and/or fever. With a few exceptions, gastroenteritis is self-limiting illness that is cleared in 5-7 days (1). *STm* can cause a successful infection by virtue of its two-distinct virulence-associated Type Three Secretion Systems, TTSS-1, and TTSS-2 encoded by *STm* pathogenicity islands 1 (SPI-1) and 2 (SPI-2), respectively. The TTSS allows the secretion of several bacterial effector proteins directly into host cytoplasm at a specified time during infection. These effectors bestow several pathogenicity related functions that include entry of *STm* into epithelial cells, its replication in hostile intracellular environments, and also reprograming of host transcriptional machinery and signaling (2). After entering into the cell, secreted factors such as SopE, SopE2, SopB along with a few others induce the activation of mitogen-activated protein kinase (MAPK) pathways leading to significant modulation of these and associated pathways. These mechanisms, which are driven by Erk, JNK and p38, eventually converge to activate the transcriptional regulators-Activator Protein-1 (AP-1) and Nuclear factor kappa light chain enhancer of activated B cells (NF-κB) (3, 4). AP-1 and NF-κB operate on their target gene promoters in a controlled and effective manner to orchestrate inflammatory diarrhea. NF-κB is a multimeric transcription factor required for activation of several target genes during infection. AP-1 is a collective term referring to dimeric transcription factors composed of members of Jun and Fos families that bind to consensus motif ‘5-TGAG/CTCA-3’ (5–7). Host epithelial pro-inflammatory signaling is a hallmark of *STm* mediated inflammatory diarrhea. Regulators such as NF-kB, STAT and AP-1 are known to get activated during the acute phase of *STm* infection initiating a pro-inflammatory environment and recruitment of neutrophils. *STm* effector SipA was shown to induce IL-8 through phosphorylation of subunit of AP-1 and p38 MAPK (8). AP-1 activity is regulated at multiple levels including-composition of the dimer, copy number, localization and stability. Post-translational modifications (PTMs) modulate each of these aspects related to AP-1 (9). While it is known that AP-1 activity is regulated by both SUMOylation and phosphorylation, roles for these PTMs in discrimination between various target genes regulation during *STm* infection is not known. Despite the activation of AP-1 expression upon *STm* infection in the nucleus only a subset of potential target genes is expressed, and the mechanisms that govern this phenomenon are yet to be explored.

Post-translational modifications (PTMs) and subsequent regulation of proteins represent an important process that is manipulated by pathogens during infection (10–12). PTMs represent a mode of rapid and reversible alteration of the proteome leading to outcomes that are detrimental to pathogenesis. Small ubiquitin like modifier (SUMO), is a Post-translational modifier known to be critical for cell signaling, transcriptional reprograming and several other fundamental cellular processes. We had reported earlier that SUMOylation pathway is down-regulated during *STm* infection (10). Specifically, simultaneous down-regulation at the transcriptional level of the gene encoding Ubc9, an E2-SUMO-conjugase enzyme and PIAS1, during *STm* infection was observed. However, the mechanism of this downregulations remained unknown. Furthermore, in another study, we showed that *STm* infection leads to an alteration of the SUMO-modified proteome (13). In this current study, we attempted to understand the relationship between SUMOylation of c-Fos and/or c-Jun and their ability to limit transcriptional activity of genes coding for SUMO pathway enzymes. Unexpectedly we observed that SUMOylation modulates differential target gene expression by c-Fos, and, in turn, the regulation of SUMOylation pathway genes, Ubc9 and PIAS1, was governed by c-Fos.

## Results

### Identification of potential AP-1 target sites in promoter regions of UBC9 and PIAS1

Simultaneous downregulation of SUMOylation pathway genes (SUMO genes) viz PIAS1 and Ubc9 (10), during *STm* infection led us to investigate the possible involvement of common regulatory mechanisms. We carried out an *in-silico* analysis of promoter region of SUMO pathway genes in an attempt to find conserved motifs that may represent binding regions of transcription factors. Using Gene Bank Database (NCBI-Gene), ∼ -10.6 Kb promoter region was analyzed, in each case, using conserved motif search (Figure 1A). Several potential regulatory motifs, including that of NF-kB, PPARγ, c-Fos, c-Jun, NFAT and CREB, could be identified in the promoter regions of UBC9 (UBE2I), PIAS1, PIAS4 and SAE1 genes (Figure 1A). Notably, we located multiple AP-1 binding motifs on the promoters of SUMO pathway genes (Figure 1B) (14–16). AP-1 regulatory motifs were significantly overrepresented and therefore were further analyzed. AP-1 is known to bind to TPA-responsive element or the AP-1 motif: ‘5-TGAG/CTCA-3’ (5–7). To reconfirm our findings by an alternative method, an online software tool Genomatix: Matinspector (17, 18) (Figure S1A) and CiiiDER (19) was also used specifically for identification of motifs for DNA binding by regulators. The analysis revealed presence of multiple potential binding motifs of Fos, Jun, Fos: Jun (heterodimer) and Jun: Jun (homodimer) in the SUMO pathway gene promoters (Figure S1B and S1C). An *in-silico* comparative alignment analysis, of AP-1 binding motif, was also done between GenBank Database (NCBI-Gene) and Genomatix: Matinspector tool (Figure 1C). The analysis confirmed presence of AP-1 regulatory motif near transcription start site (TSS, +1) of the SUMO pathway genes. Interestingly, both strategies allowed identification of the *same* consensus AP-1 binding motif in the PIAS1 promoter. Furthermore, in the UBC9 promoter, a consensus AP-1 binding motif was identified at -10.5kb using GeneBank Database. Since the matinspector software tool restricts itself to only 1kb promoter regions, we were not able to represent same -10.5kb consensus AP-1 binding motif by *in-silico* comparative alignment in UBC9 promoter. To further analyze the relative positions of AP-1 binding motif, multiple sequence alignment of promoter regions in SUMO pathway genes from transcription start site (TSS, +1) was done using EMBL-MSA: MView Program (Figure 1D). In order to validate the findings, an enrichment analysis for regulatory motif(s) was carried to identify transcription factor binding site (TFBS) that are significantly over or under represented. Specifically, the analysis focused on promoters of SUMO pathway genes and compared with promoters of background genes (Table 1) that are known not to undergo alteration upon *STm* infection (p-value p<0.05) (Figure 1E). Enrichment analysis suggested that the AP-1 binding motif is the most significant TFBS in promoters of SUMO pathway genes. Together these data allowed us to conclude that UBC9 and PIAS1 promoters harbor potential AP-1 binding motifs. Considering the role of AP-1 in the inflammation with respect to *STm* infection (20–22), we investigated its role in regulation of Ubc9 and PIAS1.

**Fig 1.**
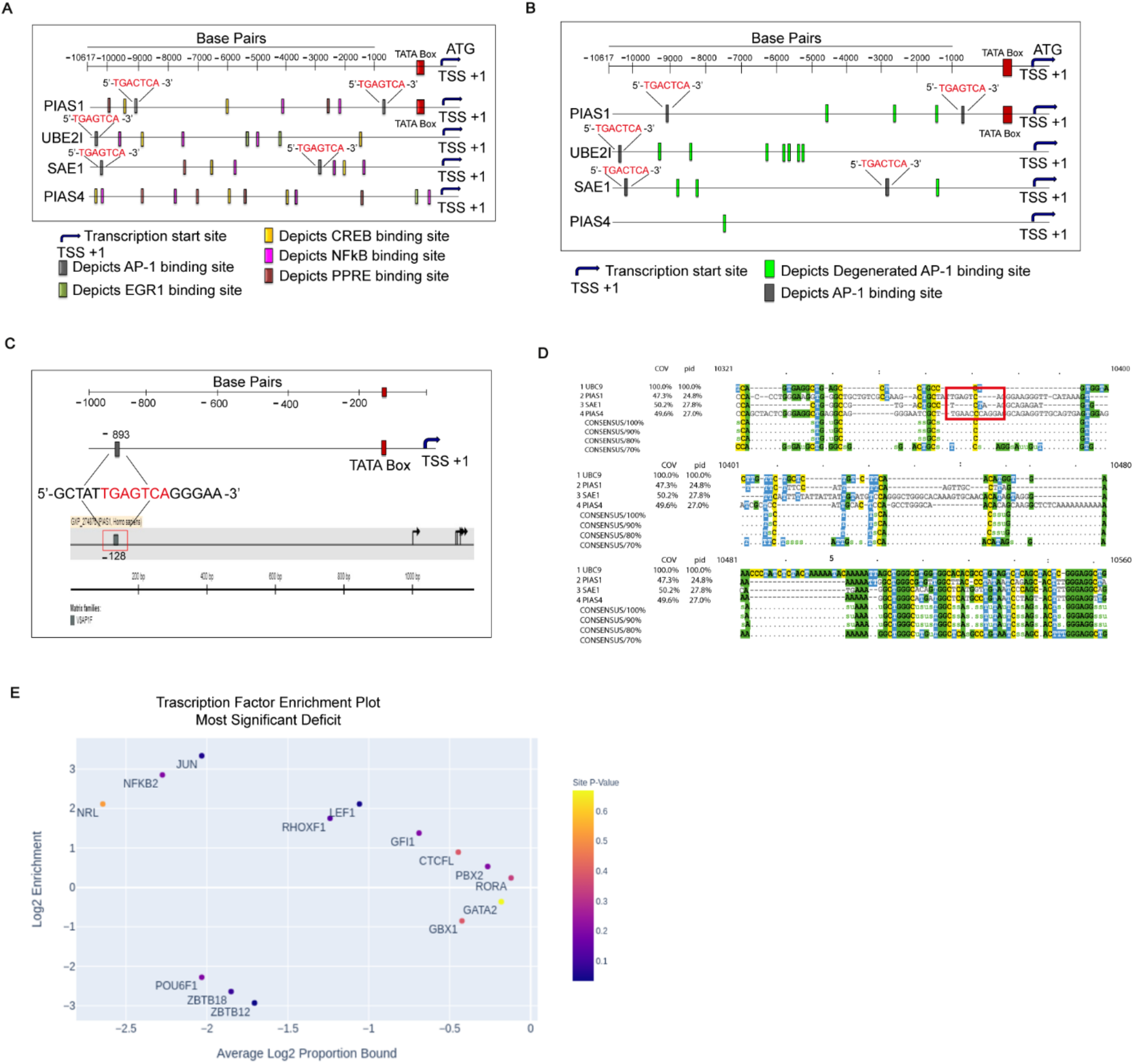
Identification of potential regulatory binding motifs by *in-silico* promoter analysis of SUMO-pathway genes. (A) *In-silico* promoter analysis was done by searching conserved motifs manually in ∼10.6 kb promoter regions of each of SUMOylation pathway genes. Promoter regions were extracted using GenBank Database (NCBI-Gene) and analyzed for consensus regulatory motifs for respective transcription factors (TFs). Analysis depicts presence of multiple consensus regulatory motifs depicted in the figure by colored rectangles in the 5’ to 3’ regions of the aligned promoters. Motifs of Activator protein (AP-1) is shown in grey rectangle. (B) Identification of several consensus and degenerated AP-1 regulatory motifs in the promoter regions of each SUMO pathway genes (indicated as grey rectangle and green box, respectively. (C) Comparative alignment of identified AP-1 regulatory motif was done between Gen Bank Database (NCBI-Gene) and Genomatix: Matinspector tool in (1kb) PIAS1 promoter. (D) Multiple sequence alignment (MSA) using MView program showing promoter of each SUMO pathway gene and represent distinct annotation along with consensus AP-1 regulatory motif. (E) Enrichment analysis showing presence of statistically significant transcription factor binding sites (TFBSs) in 5kb promoter regions of SUMO pathway genes compared to background regions (controls or unaffected genes upon *STm* infection) using JASPAR 2020 database based computational CiiiDER tool with 0.15 deficit value (core and matrix scores equal or greater than to 0.85). The plot shows enrichment (y-axis, ratio of proportion bound) and average log proportion bound (x-axis). Size and color show log10 (P-value p<0.05) scale of enrichment values greater than zero indicate over-representation of TF and less than zero indicate under-representation. Size of the circle indicate significance score.

**Table 1.** List of background genes used for enrichment analysis.

| *S. No.* | *Genes* |
| --- | --- |
| 1. | HPRT1 |
| 2. | Beta Tubulin |
| 3. | GAPDH |
| 4. | Beta Actin |
| 5. | SAE2 |

### Direct binding and regulation of SUMO genes by AP-1

Electrophoretic Mobility Shift Assay (EMSA) was used to examine possible direct physical association of c-Fos/c-Jun with the identified AP-1 regulatory motifs in PIAS1 promoter. EMSA was performed to evaluate the binding of c-Fos/c-Jun to 39 base pairs labelled fragment of PIAS1 promoter (PIAS1^PROMOTER^) that harbored the single AP-1 binding motif. A labelled fragment of CCL5 gene promoter (CCL5^PROMOTER^) was included as a positive binding control for c-Fos (23). As can be discerned from the figure 2A, presence of labeled PIAS1^PROMOTER^ and nuclear extracts of Human colonic adenocarcinoma derived cell line HCT-8 cells yielded a mobility shift band (indicated with a dark arrow, in Figure 2A, Lane2). Moreover, molar excess of unlabeled PIAS1^PROMOTER^ (100X) in the mixture diminished the PIAS1 mobility shift band (Figure 2A, Lane3). Next, we tested whether the mobility shift band was a result of binding of c-Fos by super shift assay using anti-c-Fos specific antibody. When c-Fos antibodies were added to the reaction mixture (containing PIAS1^PROMOTER^ and nuclear extracts), a super shift band was observed indicating binding of c-Fos complex to the probe (Figure 2A, Lane4). Thus, we inferred that c-Fos indeed was able to bind to the PIAS1 promoter. A PIAS1 promoter with a dinucleotide mutation in AP-1 regulatory motif (CA replaced by TG) was created (PIAS1^MUT^-^PROMOTER^). PIAS1^MUT^-^PROMOTER^ showed no change in shift band but no super shift was observed in presence of PIAS1^MUT^-^PROMOTER^ (24, 25) (Figure 2B, Lane1 & 2). However, a PIAS1 promoter completely lacking the AP-1 regulatory motif (PIAS1^NULL^-^PROMOTER^) resulted in loss of both shift and super shift bands (Figure 2B, Lane3 & 4). Together these data led us to conclude that c-Fos directly binds to ‘5-TGAGTCA-3’ motif in the PIAS1 promoter. Furthermore, it led us to hypothesize that c-Fos may be required for PIAS1 transcriptional regulation. To test this hypothesis, c-Fos/c-Jun were perturbed in HCT-8 cells that were either mock infected or infected with *STm* SL1344 strain for 4 hrs. Post infection expression analysis of SUMO genes was carried out using quantitative real time PCR (qRT-PCR). The data revealed that knocking down of c-Jun resulted in altered expression of Ubc9 and PIAS1 at mRNA and protein level compared to scrambled control (Figure 2C, S2A). Ubc9 and PIAS1 expression were downregulated when cells were knocked-down for c-Jun. The extent of down-regulation was equivalent to scrambled Si-RNA treated cells that were infected with *STm*. On the contrary, up-regulation of c-Jun by transfection of cells with c-Jun encoding plasmids resulted in up-regulation of both PIAS1 and Ubc9 (Figure 2D, S2B-C). In line with these results, we observed a positive correlation between the expression of c-Fos and Ubc9/PIAS1 (Figure 2E-F). Downregulated expression of Ubc9 and PIAS1 was seen upon c-FOS knock down compared to scrambled control. Alternatively, expression of Ubc9 and PIAS1 was up regulated in cells where c-Fos expression was enhanced by transfection of c-Fos encoding plasmid (Figure 2F, S2D). We therefore concluded that AP-1 activates the transcription of Ubc9 and PIAS1.

**Fig 2.**
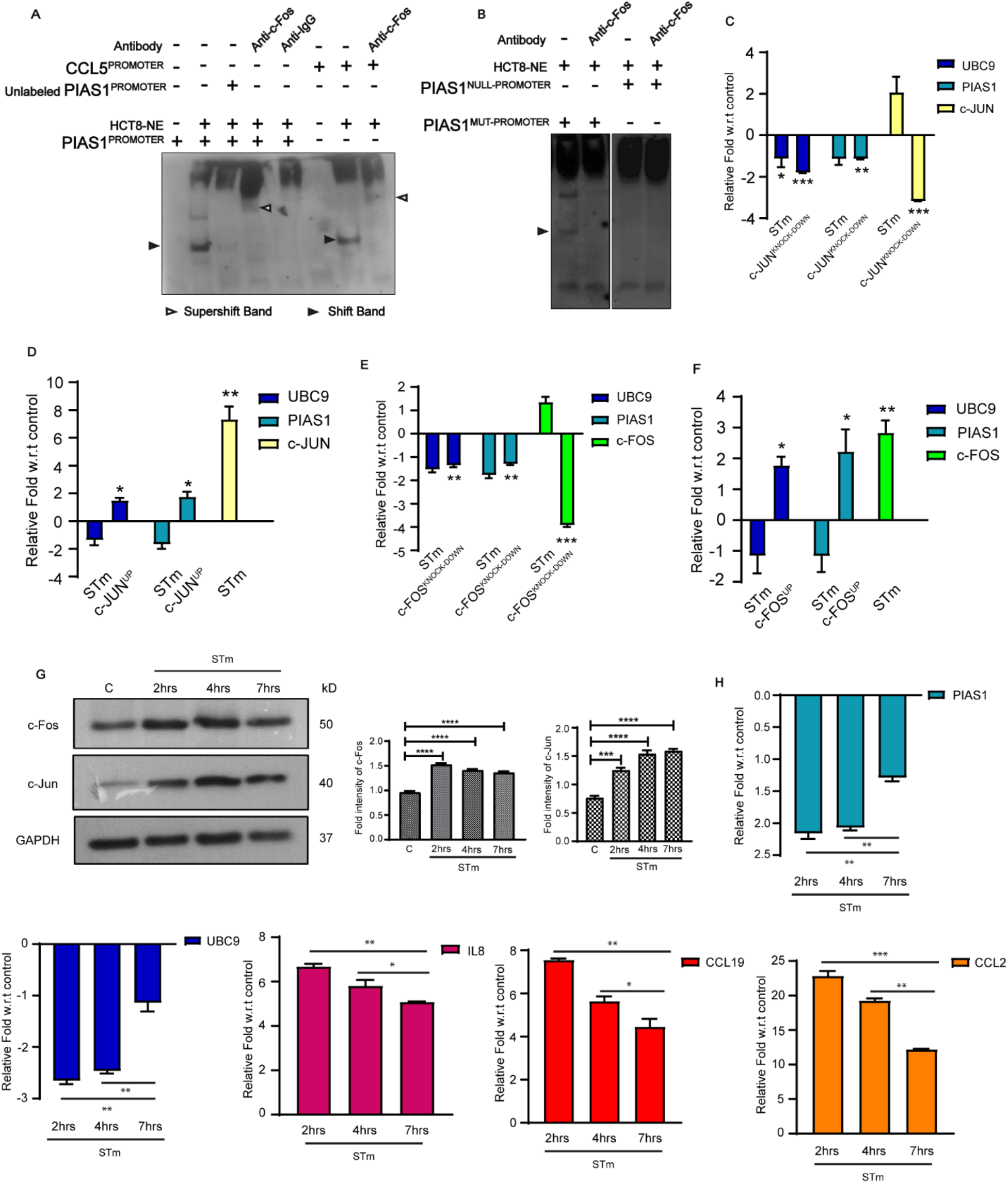
Demonstration of direct binding and regulation of SUMO gene by AP-1. (A) EMSA was carried out using labelled PIAS1 promoter harboring the putative AP-1 motif (TGAGTCA) or CCL5 promoter (as indicated) along with nuclear extracts from HCT-8 cells. Unlabeled PIAS1 promoter was used as a competitor. Addition of each reagent is shown by **+** sign. Lane 1; PIAS1 promoter only and shows no bands, Lane 2: PIAS1^PROMOTER^ was added with nuclear extract, presence of shift band marked with dark arrow, Lane 3: shows absence of ‘shift band’ in presence of (competitor) 100X unlabeled PIAS1^PROMOTER^, Lane 4, representing ‘super shift band’ (marked with open arrow) in presence of PIAS1^PROMOTER^ and nuclear extract along with anti c-Fos antibody. Equal amount of nuclear extract was loaded in each lane. (B) EMSA was carried out as described in (A) except that the PIAS1^MUT-PROMOTER^ used here harbored a mutation in AP-1 regulatory motif (CA replaced by TG), or a PIAS1^NULL-PROMOTER^ (completely lacking AP-1 regulatory motif). (C) qRT-PCR analysis showing expression of Ubc9 & PIAS1 upon Si-RNA mediated c-JUN silencing (c-JUN^KNOCK-DOWN^) and 4hrs *Salmonella* Typhimurium (*STm*) infection of HCT-8 cells compared to those treated with scrambled Si-RNA. Relative fold change compared to control untreated cells is plotted (D) qRT-PCR analysis of UBC9 and PIAS1 expression upon c-JUN encoding plasmid based over expression (c-JUN^UP^) and 4hrs *STm* infection of HCT-8 cells. Relative fold change compared to control untreated cells is plotted. (E) qRT-PCR analysis of Ubc9 & PIAS1 expression upon Si-RNA mediated c-FOS silencing (c-FOS^KNOCK-DOWN^) and 4hrs *STm* infection of HCT-8 cells compared to those treated with scrambled Si-RNA. (F) qRT-PCR analysis of Ubc9 & PIAS1 expression upon c-FOS over expression condition (c-FOS^UP^) and 4hrs *STm* of HCT-8 cells. Relative fold change compared to control is plotted. HPRT was used for normalization. Error bars represent mean of SE. Statistical analysis was done using unpaired t-test (‘***’ p-value <.001;’**’ p-value <.01; ‘*’ p-value <.05). (G) Immunoblotting for c-Fos and c-Jun during different time of *STm* infection (2hrs, 4hrs and 7hrs) in HCT-8 cells. GAPDH was used as loading control. Densitometric analysis was done using means + SE of expression data from three independent experiments for the respective protein. (H) Analysis of expression of indicated AP-1 target genes using qRT-PCR upon different time point of *STm* infection of HCT8 cells. HPRT was used for normalization. Error bars represent mean of SE. Statistical analysis was done using unpaired t-test.

In addition to SUMO pathway genes, several other target genes of AP-1 are known to be involved during *STm* infection. To understand the mechanism of their regulation by AP-1, expression dynamics of two canonical well characterized targets CCL2 and CCL19 along with Ubc9 and PIAS1 were analyzed. A time course analysis of gene expression was performed in HCT-8 cells post *STm* infection at 2hrs, 4hrs and 7hrs. Post infection (p.i), as anticipated, a gradual increase in expression of c-Fos and c-Jun was observed in response to *STm* infection (Figure 2G). However, contrary to this, the expression of Ubc9 and PIAS1 was downregulated during *STm* infection in line with our earlier reports (10, 13). Between 2hrs and 7hrs p.i, down-regulation of Ubc9 and PIAS1 was observed, while CCL2 and CCL19 were upregulated (Figure 2H). These data suggest that the target genes of AP-1 are differentially regulated upon *STm* infection.

### Differential regulation of target genes of c-Fos through Post-translational modifications

Next, we were interested to understand the basis of the differential regulation of AP-1 target genes during *STm* infection. Both c-Fos and c-Jun are known to be regulated by phosphorylation and SUMOylation (9,26–28). Using HCT-8 cells, we investigated the possible contribution of SUMOylation in selective activation of target genes, particularly in the context of *STm* infection. We restricted our investigations to c-Fos, particularly since its transcriptional activity anyway requires c-Jun as it always functions as a heterodimer (29). Control HCT-8 cells or those transfected with one of the constructs viz Wild-Type c-FOS (c-FOS^WILD-TYPE^), phospho-deficient c-FOST232A (c-FOS^PHOS-DEF^), a SUMO-deficient c-FOSK265R (c-FOS^SUMO-DEF^), were infected with *STm*. Initially, transcriptional expression of SUMO pathway genes and other AP-1 target genes was examined in these cells using qRT-PCR. These experiments revealed that unlike the cells expressing the endogenous c-Fos, overexpression of c-FOS^WILD-TYPE^ or c-FOS^SUMO-DEF^ or c-FOS^PHOS-DEF^ prevented *STm* infection-induced down regulation of Ubc9 and PIAS1 (Figure 3A). Instead a significant upregulation of Ubc9 and PIAS1 was observed. In case of CCL2 and CCL19, *STm* infection induces the expression of CCL2 and CCL19 (∼ 8-fold and 4-fold respectively). However, in cells with c-FOS^WILD-TYPE^ or c-FOS^SUMO-DEF^, a constitutive increase in expression of both CCL2 and CCL19 was seen, even without any infection. Infection led to only a marginal increase in CCL2 or CCL19 expression in cells with c-FOS^WILD-TYPE^ overexpression, and a decrease in those with c-FOS^SUMO-DEF^ overexpression (Figure 3B). Strikingly, unlike for c-FOS^SUMO-DEF^, the presence of c-FOS^PHOS-DEF^ led to downregulation of both CCL2 and CCL19 in both control and infected cells. Thus, we concluded that c-Fos SUMOylation and phosphorylation have distinct effects for different AP-1 target genes.

**Fig 3.**
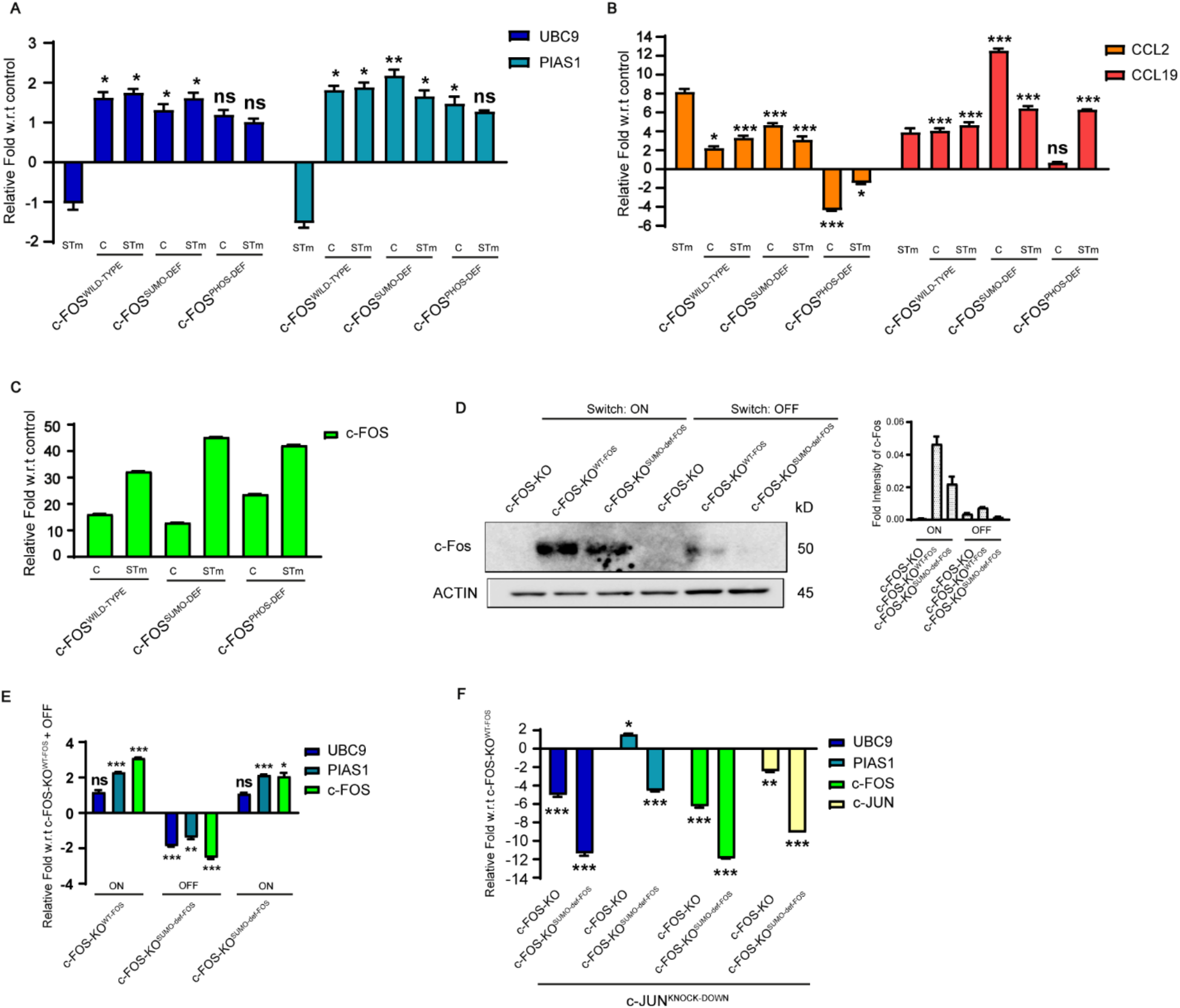
Analysis of c-Fos mediated regulation of target genes through Post-translational modifications. (A-C) c-FOS^WILD-TYPE^ and plasmids encoding different PTMs defective versions i.e. phospho-deficient c-FOST232A (c-FOS^PHOS-DEF^) and SUMO-deficient c-FOSK265R (c-FOS^SUMO-DEF^) encoding plasmids transfected HCT-8 cells which were either left untreated or infected with *STm* for 4hrs. qRT-PCR based expression analysis of SUMO genes and other AP-1 target genes was then performed. HPRT was used for normalization. (D-E) Immunoblot and qRT-PCR analysis showing expression of Ubc9, PIAS1 and c-Fos during c-Fos induction ‘ON’ and ‘OFF’ conditions in c-FOS-KO, c-FOS-KO^WT-FOS^ and c-FOS-KO^SUMO-def-FOS^ MEFs. Switch ‘ON’ represents exposure of these cells to 20% serum containing DMEM media, while Switch ‘OFF’ condition indicates absence of these cell to 20% serum containing DMEM media. (F) qRT-PCR analysis of SUMO with AP-1 target genes from c-FOS-KO and c-FOS-KO^SUMO-def-FOS^ compared to scrambled c-FOS-KO^WT-FOS^ upon c-JUN^KNOCK-DOWN^ condition. Actin was used as loading control. Densitometric analysis of expression by means + SEM from three independent experiments for the respective protein. Error bars represent SE. Statistical analysis was done using unpaired t-test and one-way ANOVA (multiple comparisons).

To negate the effect of endogenous c-Fos on the outcome of these experiments, we carried our expression analysis in mouse embryonic fibroblasts (MEFs) line with stable disruption of endogenous c-FOS (by homologous recombination) (hereafter to be called as c-FOS-KO cells) along with those expressing wild type c-FOS (c-FOS-KO^WT-FOS^) or SUMO deficient c-FOSK265R (c-FOS-KO^SUMO-def-FOS^) under a Serum Response Element (27,30,31). These cells were cultured under serum deprivation for 16hrs followed by addition of high serum (20%) containing DMEM media to induce the expression of the respective forms of c-Fos during switch ‘ON’ condition (27). Addition of serum in the media allowed significant induction of c-Fos protein expression (Figure 3D). MEFs expressing c-FOS-KO^SUMO-Def-FOS^ and those expressing c-FOS-KO^WT-FOS^ showed upregulation of expression of Ubc9 and PIAS1 at RNA level compared to cells without c-Fos expression (Figure 3E). In the case of c-FOS-KO^SUMO-Def-FOS^ in “OFF’ condition, a downregulation of Ubc9 and PIAS1 was observed at RNA level. Surprisingly, at protein level we observed higher expression of Ubc9 and PIAS1 during “OFF’ condition (absence of c-Fos expression) (Figure S3A), indicating dysregulation of SUMO-gene expression due to the absence of c-Fos. To examine if increase in expression of Ubc9 and PIAS1 during *STm* infection in c-FOS-KO cells was due to c-Jun/c-Jun dimer activity, c-Jun was knocked down in these cells using specific Si-RNA. The c-Jun Si-RNA treatment led to a drastic reduction in Ubc9 and PIAS1 expression (Figure 3F), indicating that in the absence of c-Fos, homodimers of c-Jun-c-Jun drives the expression of Ubc9 and PIAS1. Next, we performed EMSA using Murine c-FOS-KO MEFs. Nuclear lysates from c-FOS-KO MEFs when added to the EMSA reaction mix revealed the binding of PIAS1^PROMOTER^ with nuclear extracted proteins of c-FOS-KO MEFs. As shown in Figure S3C, Lane2, a mobility shift is seen when the DNA was mixed with lysates. This shift may be due to presence of c-Jun or other proteins belonging to AP-1 family. Here, we conclude that AP-1 binds to the identified AP-1 regulatory motif in PIAS1 promoter using c-FOS-KO MEFs. These c-FOS-KO MEFs do not have c-Fos protein, which explain the absence of super shifted band (Figure S3C, Lane 4). Together, these data reveal that c-Fos and c-Jun regulate target genes in a coordinated manner, the PTM variants c-FOS-KO^SUMO-Def-FOS^ and c-FOS-KO^WT-FOS^ display target gene activation in a manner which is distinct. Thus, localization of transcription factor in the nucleus alone is not the only requirement for target gene activation. Together, these results indicated that PTM modification of c-Fos has a distinct effect on transcription of its target genes.

### c-Fos SUMOylation mediates immune gene activation and controls inflammation

Next, we investigated possible physiological cellular consequences of differential regulation of target genes in host during *STm* infection on the cellular transcriptome. Murine MEFs line c-FOS-KO^WT-FOS^ and c-FOS-KO^SUMO-def-FOS^ (described in the previous section) were used (Figure 4A). *STm* infections were carried out in c-FOS-KO^WT-FOS^ and c-FOS-KO^SUMO-def-FOS^ MEFs cells for 4hrs during switch ‘ON’ condition, since this time corresponded to the maximal c-Fos activity expression. Total RNA from these samples were extracted and subjected to 3’mRNA sequencing (Figure 4A). Multiple pairwise comparisons were done to identify differentially regulated genes (Figure 4B). A total of 2416 genes were differentially regulated with an FDR < 0.05 combined in all the comparisons (Figure S4B). The differentially expressed genes were further represented as a Venn diagram to emphasize on the number of genes unique to a specific comparison and common between two or more comparisons (Figure 4C). No changes were observed when mock-treated c-FOS-KO^WT-FOS^ [c-FOS-KO^WT-FOS^ (UI-uninfected)] were compared with mock-treated c-FOS-KO^SUMO-def-FOS^ cells [c-FOS-KO^SUMO-def-FOS^ (UI)] (Figure 4B), indicating that during steady-state, SUMOylation of c-Fos may not have a significant role in the control of gene expression. Interestingly, the comparison between *STm* infected c-FOS-KO^SUMO-def-FOS^ [c-FOS-KO^SUMO-def-FOS^ (I)] versus *STm* infected c-FOS-KO^WT-FOS^ [c-FOS-KO^WT-FOS^ (I-infected)] showed a total of 1534 dysregulated genes, 793 genes being upregulated, and 741 down-regulated (Figure 4B and 4F). Gene ontology (GO) analysis on 2416 genes (that were significantly differentially expressed in the three pairwise comparisons combined) revealed that GO terms like nucleic acid binding, response to stress, response to cytokine, TNF signaling pathway were significantly enriched (Figure S4D). Biological process analysis in c-FOS-KO^SUMO-def-FOS^ (I) versus c-FOS-KO^WT-FOS^ (I) samples suggested that pathways related to Cell adhesion, Cell migration, Response to growth factor, Regulation of signal transduction and cell communication were downregulated while pathways related to Immune system, Defense response, Metabolic process, Response to cytokine etc. were upregulated suggesting that components of key pathways are differentially regulated in c-FOS-KO^SUMO-def-FOS^ versus c-FOS-KO^WT-FOS^ post-infection (Figure 4G). We further looked at the expression patterns of a set of genes associated with the affected pathways during *STm* infection and subsequent inflammation (Figure 4H). Subset of these dysregulated genes included-those having a known role in *STm* infections, immune response pathway genes and those involved in defense response (oxidative stress) etc. Representative examples of these genes are listed in (Table 6). As shown in the heat map (Table 2), several immune regulatory genes were found to be significantly upregulated, including AP-1 target gene CCL2, CXCL1, in c-FOS-KO^SUMO-def-FOS^ (I) compared to c-FOS-KO^WT-FOS^ (I) samples (Figure 4H). According to transcriptome analysis CCL2 and CXCL1 expression was higher in c-FOS-KO^SUMO-def-FOS^ (I) compared to c-FOS-KO^WT-FOS^ (I) samples. For validation, a few immune genes and SUMO pathway genes were picked and were analyzed using qPCR (Figure S4C). gProfiler was used to generate KEGG pathway graphs. To investigate the affected biological pathway, gene ontology analysis was done using pairwise comparison of DEGs between c-FOS-KO^SUMO-def-FOS^ (I) and c-FOS-KO^WT-FOS^ (I) samples (Figure S4D) (32). In line with the above results, Gene Ontology analysis revealed that several of the upregulated genes were associated to pathways related to critical cellular processes including immune response regulation (Figure 4G) (33). These data highlight the importance of the SUMOylation of c-Fos in control of immune responses during *STm* infection.

**Fig 4.**
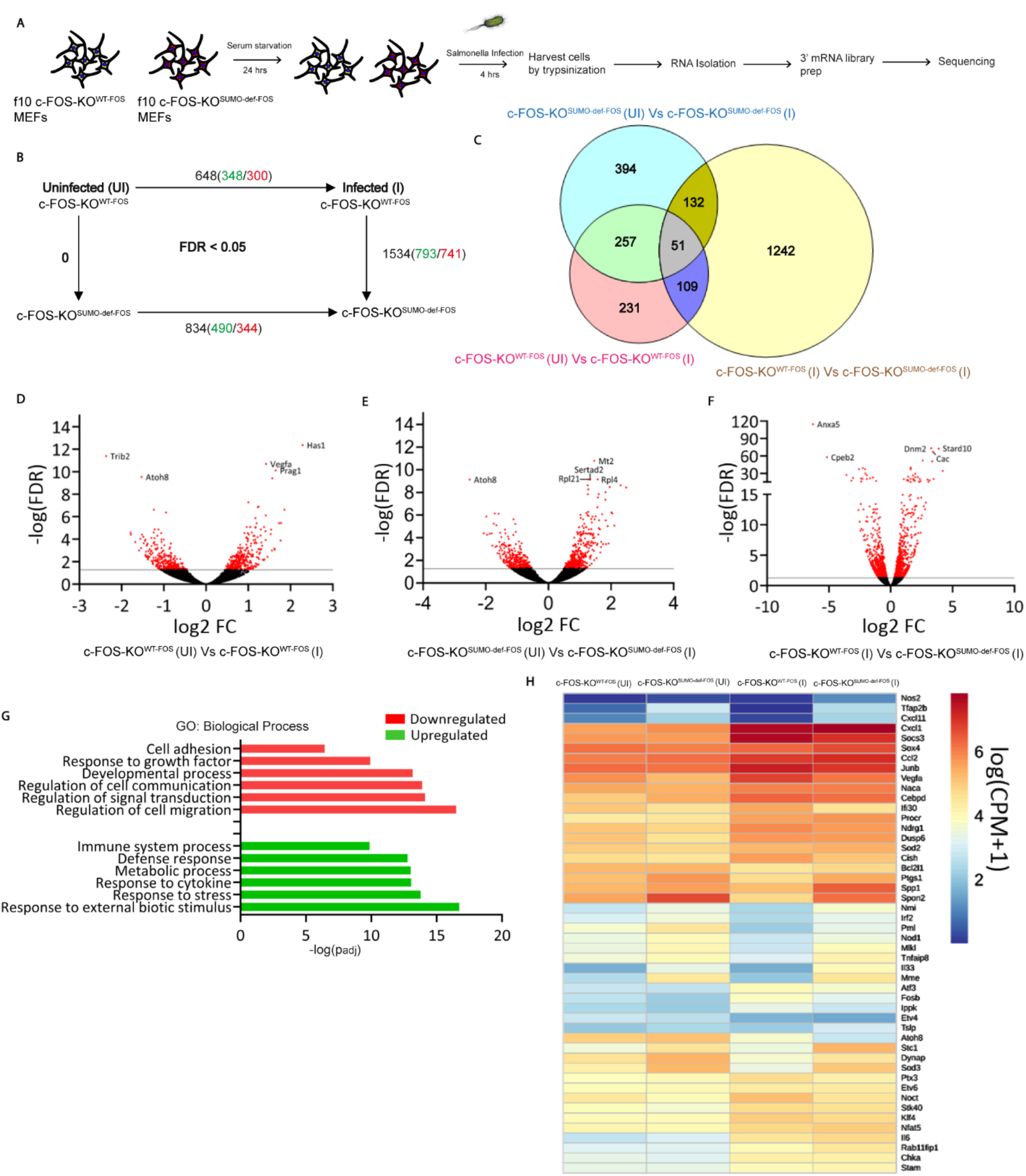
mRNA-seq analysis identifies c-Fos PTM mediated modulation of immune response in MEFs upon *STm* infection. (A) Schematic representation workflow of 3’ mRNA sequencing performed at three independent replicates. (B) Schematic representing the pairwise comparison of two experimental conditions. Numbers shown in black represent the total significantly differentially expressed genes in each pairwise comparison. Numbers shown in red represent downregulated genes and numbers shown in green represent upregulated genes. (UI)-Uninfected, (I) *STm* infected at 4hrs. (C) Venn diagram representing the overlap of number of differentially expressed genes among different pairwise comparisons. Number of genes unique to a specific set are also represented. (D-F) Volcano plots showing the genes altered in different pairwise comparisons. Genes that are significantly differentially expressed (FDR<0.05) are highlighted in red. Log 2-fold change values are plotted on the X-axis and the -log(FDR) plotted on the Y-axis. The top five most significantly differentially expressed genes are shown. (G) GO representation of Biological process analysis that are significantly enriched in c-FOS-KO^SUMO-def-FOS^ (I) versus c-FOS-KO^WT-FOS^ (I). (H) Heat map representing the normalized expression counts of a key subsets of genes.

**Table 2.** List of differentially expressed genes (DEGs) upon pairwise comparison.

| *ID* | *Gene Symbol* |
| --- | --- |
| ENSMUSG00000031838 | IFI30 |
| ENSMUSG00000024512 | DYNAP |
| ENSMUSG00000014813 | STC1 |
| ENSMUSG00000023951 | VEGFA |
| ENSMUSG00000042608 | STK40 |
| ENSMUSG00000036986 | PML |
| ENSMUSG00000027611 | PROCR |
| ENSMUSG00000024843 | CHKA |
| ENSMUSG00000019960 | DUSP6 |
| ENSMUSG00000071637 | CEBPD |
| ENSMUSG00000053113 | SOCS3 |
| ENSMUSG00000037621 | ATOH8 |
| ENSMUSG00000027820 | MME |
| ENSMUSG00000024810 | IL33 |
| ENSMUSG00000006818 | SOD2 |
| ENSMUSG00000072941 | SOD3 |
| ENSMUSG00000037379 | SPON2 |
| ENSMUSG00000025927 | TFAP2B |
| ENSMUSG00000026946 | NMI |
| ENSMUSG00000012519 | MLKL |
| ENSMUSG00000029304 | SPP1 |
| ENSMUSG00000029380 | CXCL1 |
| ENSMUSG00000031488 | RAB11FIP1 |
| ENSMUSG00000026718 | STAM |
| ENSMUSG00000076431 | SOX4 |
| ENSMUSG00000061315 | NACA |
| ENSMUSG00000007659 | BCL2L1 |
| ENSMUSG00000003847 | NFAT5 |
| ENSMUSG00000023087 | NOCT |
| ENSMUSG00000032578 | CISH |
| ENSMUSG00000003032 | KLF4 |
| ENSMUSG00000021385 | IPPK |
| ENSMUSG00000027832 | PTX3 |
| ENSMUSG00000017724 | ETV4 |
| ENSMUSG00000030199 | ETV6 |
| ENSMUSG00000005125 | NDRG1 |
| ENSMUSG00000060183 | CXCL11 |
| ENSMUSG00000003545 | FOSB |
| ENSMUSG00000052837 | JUNB |
| ENSMUSG00000025746 | IL6 |
| ENSMUSG00000035385 | CCL2 |
| ENSMUSG00000020826 | NOS2 |
| ENSMUSG00000038058 | NOD1 |
| ENSMUSG00000032487 | PTGS2 |
| ENSMUSG00000062210 | TNFAIP8 |
| ENSMUSG00000024379 | TSLP |
| ENSMUSG00000031627 | IRF2 |

**Table 3:** List of differentially expressed genes (DEGs) across the categories. The DEGs are obtained by taking a fold change cut off of 1.

| *Categories* | *Upregulated*<br>*DEGs* | *Downregulated*<br>*DEGs* |
| --- | --- | --- |
| c-FOS-KO<sup>SUMO-def-FOS</sup> (UI) vs c-FOS-KO<sup>SUMO-def-FOS</sup> (I) | 5259 | 5674 |
| c-FOS-KO<sup>WT-FOS</sup> (I) vs c-FOS-KO<sup>SUMO-def-FOS</sup> (I) | 6088 | 6412 |
| c-FOS-KO<sup>WT-FOS</sup> (UI) vs c-FOS-KO<sup>WT-FOS</sup> (I) | 6983 | 6758 |

**Table 4:** The order and size of the directional network of each category. The first column denotes the total number of proteins. the second column denotes the nodes additionally acquired (termed as mediators) from Signor database to construct a connected network.

| *categories* | *Initial total proteins* | *Mediators* | *Network Order* | *Network size* |
| --- | --- | --- | --- | --- |
| c-FOS-KO<sup>SUMO-def-FOS</sup> (UI) vs c-FOS-KO<sup>SUMO-def-FOS</sup> (I) | 13741 | 65 | 418 | 578 |
| c-FOS-KO<sup>WT-FOS</sup> (I) vs c-FOS-KO<sup>SUMO-def-FOS</sup> (I) | 12500 | 58 | 448 | 615 |
| c-FOS-KO<sup>WT-FOS</sup> (UI) vs c-FOS-KO<sup>WT-FOS</sup> (I) | 10933 | 52 | 492 | 681 |

**Table 5:** Critical driver nodes and the hubs proteins among each category.

| *Category* | c-FOS-KO<sup>WT-FOS</sup> (I)<br>vs c-FOS-KO<sup>SUMO-def-FOS</sup> (I) | c-FOS-KO<sup>WT-FOS</sup> (UI)<br>vs c-FOS-KO<sup>WT-FOS</sup> (I) | c-FOS-KO<sup>SUMO-def-FOS</sup> (UI) vs c-FOS-KO<sup>SUMO-def-FOS</sup> (I) |
| --- | --- | --- | --- |
| Numbers | 135 | 147 | 122 |
| Hub proteins | AKT2<br>MECP2<br>PRKACA<br>PRKCA<br>PTPRC<br>SIRT1<br>SMO<br>TLN1 | DMTF1<br>MECP2<br>PRKACA<br>PRKCA<br>PTPRC<br>SIRT1<br>SMO<br>TLN1 | MECP2<br>PRKACA<br>PRKCA<br>PTPRC<br>SIRT1<br>SMO |

**Table 6:** List of identified dysregulated genes having a defined role in STm infections, immune response pathway genes and defense response.

| *Genes* | *Functions* | *References* |
| --- | --- | --- |
| NOS2 | Plays a role in iron homoeostasis, generation of inflammatory chemokines and cytokines accompanied to reduce bacterial numbers in host | (59–61) |
| CXCL11 | Associated with higher bacteria burden in ceacum due to compromised intestinal IgA content, CD4<sup>+</sup> T cells and neutrophils recruitment upon 30-45 days of *STm* infection | (62,63) |
| CCL2 | Recruitment of inflammatory macrophages at colitis tissue and controls immune response and *STm* survival | (64,65) |
| IRF2 | Maturation and development of several immune cells. Display non canonical inflammasome response upon gram negative bacterial infection | (67,68) |
| IL33 | Regulator of gut permeability. Generates antimicrobial defense upon *STm* infection | (83) |
| IL6 | Modulates innate and adaptive immune response during infection and tissue injury | (21) |
| VEGF | Upon *STm* infection exerts as anti-angiogenic therapy | (84,85) |
| TNFAIP8 | Regulates host apoptosis process and pathogen invasion during *STm* infection | (86,87) |
| PTGS1 | Regulates inflammatory response of host via neutrophils chemotaxis during *STm* infection | (69–71) |
| CXCL1 | Associated with neutrophils recruitment activation and bacterial clearance | (88) |

### ***c***-Fos SUMOylation and STm mediated differential activation of host immune genes alter signaling and intracellular STm survival

Next, we have used protein-protein interaction (PPI) network analysis to analyze the RNAseq data. A total of 5259 up-and 5674 downregulated genes were identified (with a cutoff of 1 fold) in c-FOS-KO^SUMO-def-FOS^ (I) compared to c-FOS-KO^SUMO-def-FOS^ (UI). Furthermore, 6088 genes were up regulated and 6412 genes were down regulated in c-FOS-KO^SUMO-def-FOS^ (I) compared to c-FOS-KO^WT-FOS^ (I). A summary of the differentially expressed genes is shown in (Table 3). These DEGs are then used to generate directional PPI networks using signor database (Table 4). The network edges are provided in the file **network.xlsx.** Network analysis shows all three categories had a upstream protein of c-Fos like cold shock domain containing E1 (CSDE1) is that drives signal to c-Fos and has role in translational reprogramming and diseases (34). CSDE1 is a cytoplasmic RNA binding protein that controls abundance, translation of RNAs and drives post transcriptional program to promote melanoma invasion and metastasis (35, 36). Several lines of evidence have established the role of CSDE1 in cancer and inflammatory diseases (37). The networks for each category are showed in (Figure S5A-C). PPI networks can be used to tailor the complex interactions between proteins, and their efficient analysis can reveal the core group driving the system’s perturbations (38). To identify the core set of proteins, we opted for structural controllability analysis that searches for a minimal number of driver nodes that are sufficient to control the network, i.e., changes in the state of these nodes are sufficient to drive the system from any initial to any final states. The driver nodes set (MDS) in a network is not unique, but the cardinality of MDS remains the same. We found that 67.19%, 69.31%, and 66.03% nodes are driver nodes in the “c-FOS-KO^WT-FOS^ (I) vs c-FOS-KO^SUMO-def-FOS^ (I)”, “c-FOS-KO^WT-FOS^ (UI) vs c-FOS-KO^WT-FOS^ (I)” and “c-FOS-KO^SUMO-def-FOS^ (UI) vs c-FOS-KO^SUMO-def-FOS^ (I)” respectively. A larger number of driver nodes are appearing because of the sparse nature of the networks. The results of the control theory analysis are shown in (Figure 5A). By simulating the system 1000 times, we obtained 1000 different maximal matching sets. Using a simple set theory approach, we identified the Critical Drive Nodes (CDNs) in each category (Table 5 and file **critical driver nodes.xlsx**). We found that 30.13%, 29.88%, and 29.19% number of nodes were CDN in the “c-FOS-KO^WT-FOS^ (I) vs c-FOS-KO^SUMO-def-FOS^ (I)”, “c-FOS-KO^WT-FOS^ (UI) vs c-FOS-KO^WT-FOS^ (I)” and “c-FOS-KO^SUMO-def-FOS^ (UI) vs c-FOS-KO^SUMO-def-FOS^ (I)” respectively. We also observed a high degree of overlap between the CDNs of each category (Figure 5B). The pathways related to all CDNs are provided in the file **enrichment.xlsx** (Figure 5C). Interestingly, we observed that many pathways were commonly affected in each category. Therefore, we looked for unique affected pathways related to each category. mTOR signaling pathway, glioma and chronic myeloid leukemia were uniquely affected in c-FOS-KO^SUMO-def-FOS^ (UI) vs c-FOS-KO^SUMO-def-FOS^ (I) compared to other two categories while Chagas disease and HIF-1 signaling pathway were uniquely altered in c-FOS-KO^WT-FOS^ (UI) vs c-FOS-KO^WT-FOS^ (I) compared to other two categories. Although, VEGF and Rap1 signaling pathway were solely modulated in c-FOS-KO^WT-FOS^ (I) vs c-FOS-KO^SUMO-def-FOS^ (I) compared to other two categories. Moreover, Th1 & Th2 cell differentiation and T cell receptor signaling pathway were affected in both c-FOS-KO^WT-FOS^ (UI) vs c-FOS-KO^WT-FOS^ (I) and c-FOS^WT-FOS^ (I) vs c-FOS^SUMO-def-FOS^ (I) category (Figure 5C). Also, the investigation of the local properties of the network revealed that 8, 8, and 6 CDNs were hubs in the “c-FOS-KO^WT-FOS^ (I) vs c-FOS-KO^SUMO-def-FOS^ (I)”, “c-FOS-KO^WT-FOS^ (UI) vs c-FOS-KO^WT-FOS^ (I)” and “c-FOS-KO^SUMO-def-FOS^ (UI) vs c-FOS-KO^SUMO-def-FOS^ (I)” respectively. The names of these hub-CDN proteins are shown in Table 5. The fewer hubs among the CDNs justify the fact that the driver nodes tend to avoid the high degree nodes. The crucial KEGG pathways related with the hub-CDNs are shown in (Figure S6A-C). Hub-CDNs are mainly associated with cell proliferation and migration signaling pathways in all three categories. Further, we also explored the significance of uniquely identified CDNs-proteins represented by Venn diagram in each category upon gene ontology analysis. We identified that 3, 14, and 1 number of nodes were uniquely presented in the “c-FOS-KO^WT-FOS^ (I) vs c-FOS-KO^SUMO-def-FOS^ (I)”, “c-FOS-KO^WT-FOS^ (UI) vs c-FOS-KO^WT-FOS^ (I)” and “c-FOS-KO^SUMO-def-FOS^ (UI) vs c-FOS-KO^SUMO-def-FOS^ (I)” respectively. The panther gene analysis related to the uniquely identified CDNs-proteins are provided in (Figure 5D). Pathway analysis in case of c-FOS-KO^SUMO-def-FOS^ (UI) vs c-FOS-KO^SUMO-def-FOS^ (I) couldn’t be performed because only one unique CDNs-proteins was observed. Notably, many biological and immune related pathways were affected including Interleukin signaling, PI3 kinase pathway, Hypoxia response, endothelin signaling pathway etc. in c-FOS-KO^WT-FOS^ (I) vs c-FOS-KO^SUMO-def-FOS^ (I) category. However, B cell activation, Interferon gamma signaling, oxidative stress response, Ras signaling pathway etc. were modulated pathways in c-FOS-KO^WT-FOS^ (UI) vs c-FOS-KO^WT-FOS^ (I). Although, angiogenesis, inflammation mediated chemokine and cytokine signaling, apoptosis, PDGF signaling, EGF receptor signaling etc. were commonly affected in both c-FOS-KO^WT-FOS^ (I) vs c-FOS-KO^SUMO-def-FOS^ (I) and c-FOS-KO^WT-FOS^ (UI) vs c-FOS-KO^WT-FOS^ (I) category. Together these results indicate that c-Fos transcriptionally regulates variety of genes but SUMOylation of c-Fos drives the transcriptional reprogramming of target genes and alters the host signaling and immune response in context of *STm* infection.

**Fig 5.**
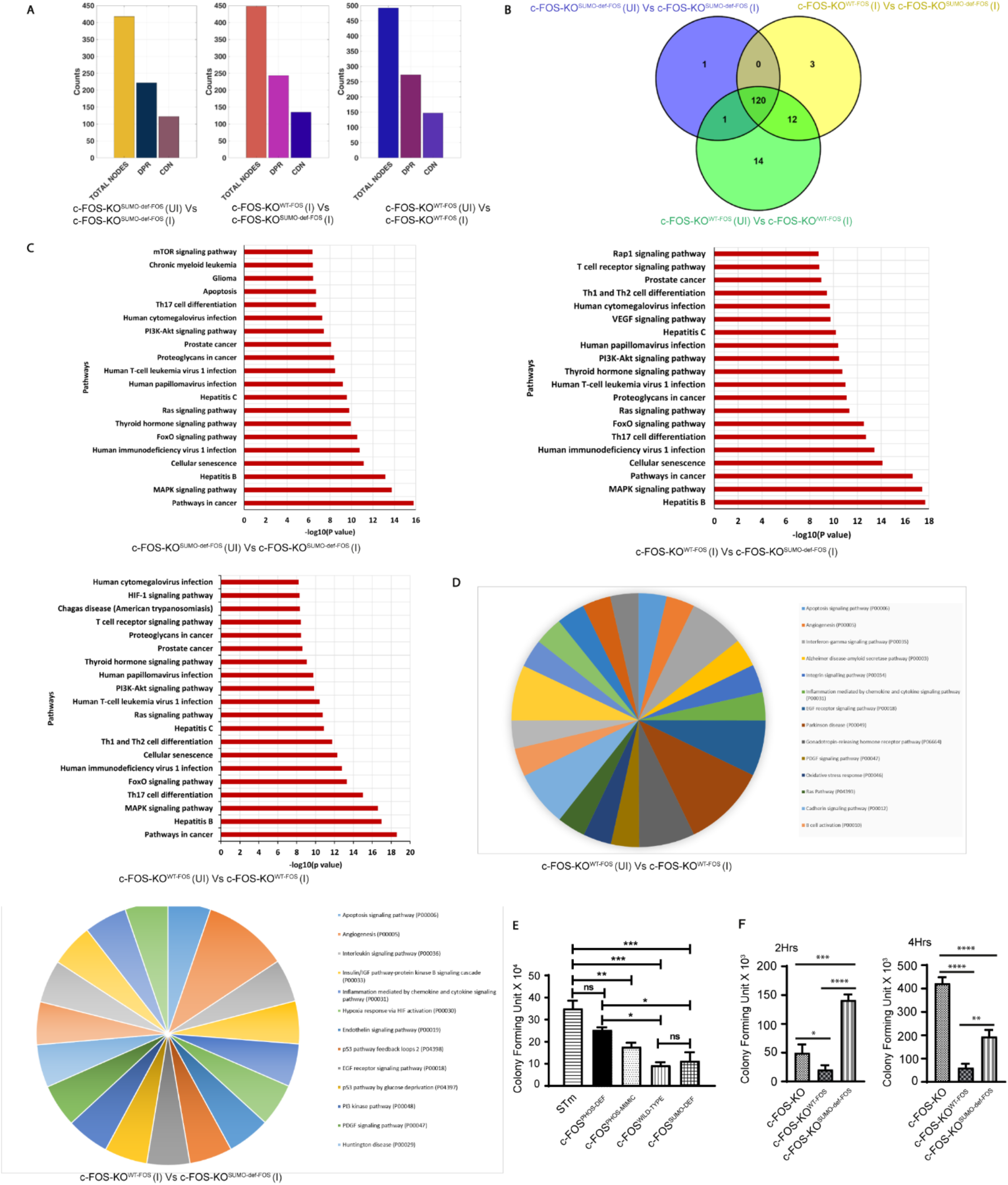
c-Fos SUMOylation and *STm* mediated differential activation of host immune response. (A) Depicts control theory analysis for each category (as mention previously) in c-FOS-KO^WT-FOS^ and c-FOS-KO^SUMO-def-FOS^ MEFs. Here, DPR: Driver nodes per run and it denotes the cardinality of the maximum matching set. CDN: Critical driver nodes in the network. (B) Venn diagram of the CDNs of the three categories revealed a high degree of overlapping between the CDNs. (C) The crucial pathways related to all CDNs of the three pairwise comparison. (D) The panther gene analysis related to the uniquely identified CDNs-proteins from c-FOS-KO^WT-FOS^ (UI) vs c-FOS-KO^WT-FOS^ (I) and c-FOS-KO^WT-FOS^ (I) vs c-FOS-KO^SUMO-def-FOS^ (I). (E) Burden assay at 4hrs of *STm* infection in transfected HCT-8 cells with c-FOS^PHOS-DEF^, phospho-mimic of c-FOST232D (c-FOS^PHOS-MIMIC^), c-FOS^WILD-TYPE^ and c-FOS^SUMO-DEF^ encoding plasmids. (F) Burden Assay at 2hrs and 4hrs of *STm* infection in c-FOS-KO, c-FOS-KO^WT-FOS^ and c-FOS^SUMO-DEF^ MEFs. HPRT was used for normalization. Error bars represent SE. Statistical analysis was done using Student’s t-test and one-way ANOVA.

Next, to test the involvement of AP-1 and its PTMs on multiplication of intracellular *STm* in intestinal epithelial cells, gentamycin protection assay (GPA) was carried out in HCT-8 cells. Healthy and *STm* infected HCT-8 cells overexpressing c-FOS^WILD-TYPE^ and different mutants of PTMs of c-FOS (c-FOS^PHOS-DEF^, phosphor-mimetic c-FOST232D (c-FOS^PHOS-MIMIC^), c-FOS^SUMO-DEF^) were tested. At 4hrs p.i. a significant lowering of *STm* colony forming units (CFU) was observed in c-FOS^WILD-TYPE^ expressing cells (Figure 5E). c-FOS^PHOS-DEF^ and c-FOS^PHOS-MIMIC^ were less efficient than c-FOS^WILD-TYPE^ at decreasing CFU and no significant differences were observed between c-FOS^SUMO-DEF^ and c-FOS^WILD-TYPE^ (Figure 5E). To investigate more accurately the consequence of c-Fos SUMOylation on the multiplication of intracellular *STm,* GPA was performed at early time (2hrs) of *STm* infection in the c-FOS-KO, c-FOS-KO^WT-FOS^, c-FOS-KO^SUMO-def-FOS^ MEFs. At 2hrs p.i, CFU was higher in c-FOS-KO^SUMO-def-FOS^ compared to c-FOS-KO and c-FOS-KO^WT-FOS^ MEFs. Surprisingly, 4hrs post *STm* infection, we observed higher number of CFU in c-FOS-KO compared to other cells. However, CFU remained higher in c-FOS-KO^SUMO-def-FOS^ compared to c-FOS-KO^WT-FOS^ MEFs (Figure 5F). The differences were not due to any cell death as apoptosis marker caspase-3 remained unchanged between c-FOS-KO, c-FOS-KO^WT-FOS^, c-FOS-KO^SUMO-Def-FOS^ MEFs at 4hrs post *STm* infection (Figure S5D). Taken together, our data suggest that AP-1, through the transcriptional regulation of genes leading to altered inflammatory environment can limit the intracellular survival of *STm*. Moreover, SUMOylation increases c-Fos ability to restrict *STm* survival

### SUMOylation dependent STm affects sub-cellular localization of c-Fos

Next, possible alteration in sub-cellular localization of c-Fos as a result of the status of its PTM modification was examined. First, we analyzed the sub-cellular localization of endogenous c-Fos in control and *STm* infected for 4hrs p.i using subcellular fractionation (39). A slight increase in the total levels of c-Fos was observed upon *STm* infection in the whole cell extract. Infection with *STm* triggered a massive increase in the nuclear localization of c-Fos and only a mild increase in the cytoplasmic pools denoting an infection mediated re-localization into the nucleus (Figure 6A). These results were confirmed by imaging experiments using HCT-8 cells infected with mCherry labelled *STm* and stained for c-Fos (Figure S7A). Label distribution graphs showed fluorescence intensity of c-Fos and Nucleus (Hoechst) along cell axis (Figure 6B). Similar to the fractionation experiments, these data pointed towards cytoplasmic to nuclear re-localization of c-Fos during infection (Figure 6B). SUMOylation of c-Fos/c-Jun was shown to limit AP-1 transcriptional activation (9,27,28,40) while its phosphorylation was shown to activate the transcriptional activity of AP-1 (41–44). To investigate whether c-Fos PTMs alter its localization, we carried out sub-cellular fractionation of mock treated and *STm* infected HCT-8 cells that were transfected with plasmids encoding either a c-FOS^WILD-TYPE^, c-FOS^PHOS-DEF^, c-FOS^SUMO-DEF^ and c-FOS^PHOS-MIMIC^. Presence of c-Fos in control and infected cells was examined in each of the fractions viz cytoplasmic, nuclear and whole cell by immunoblotting. In c-FOS^WILD-TYPE^ expressing cells, nuclear fraction of c-Fos protein showed an increase, while the cytoplasmic pool c-Fos showed a significant decrease during *STm* infection (Figure 6C). Both nuclear and cytoplasmic fractions from c-FOS^PHOS-DEF^ expressing cells showed a higher fraction of c-Fos in the nucleus in mock treated and during *STm* infection. Moreover, the fraction of cytoplasmic c-Fos showed a significant increase during *STm* infection. Compared to c-FOS^WILD-TYPE^, the c-FOS^PHOS-DEF^ showed a dramatic increase upon *STm* infection in all the three fractions (Figure 6D). Similar results were obtained with c-FOS^PHOS-MIMIC^ (Figure 6E). Cells expressing c-FOS^SUMO-DEF^ showed presence of significant amount of the protein in the nucleus even without *STm* infection and low cytoplasmic pools (Figure 6F). Moreover, limited changes in the distribution of c-FOS^SUMO-DEF^ were observed between mock-treated versus *STm* infected cells (Figure 6F). Confocal microscopy confirmed re-localization of c-FOS^WILD-TYPE^ during infection. However, in line with the above data, the SUMO deficient c-FOS^SUMO-DEF^ was localized in the nucleus even without *STm* infection (Figure S7B). Finally, we validated altered sub cellular localized expression of c-Fos from mock-treated and 4hrs *STm* infected lysates of c-FOS-KO^WT-FOS^, c-FOS-KO^SUMO-Def-FOS^ MEFs upon sub cellular fractionation by using whole cell, cytosolic and nuclear extracted lysates (Figure 6G). Together these data suggest that c-Fos PTMs alter its subcellular distribution. Based on this, we hypothesized that the differential PTM which led to alterations in sub-cellular localization c-Fos may contribute to selective regulation target gene expression.

**Fig 6.**
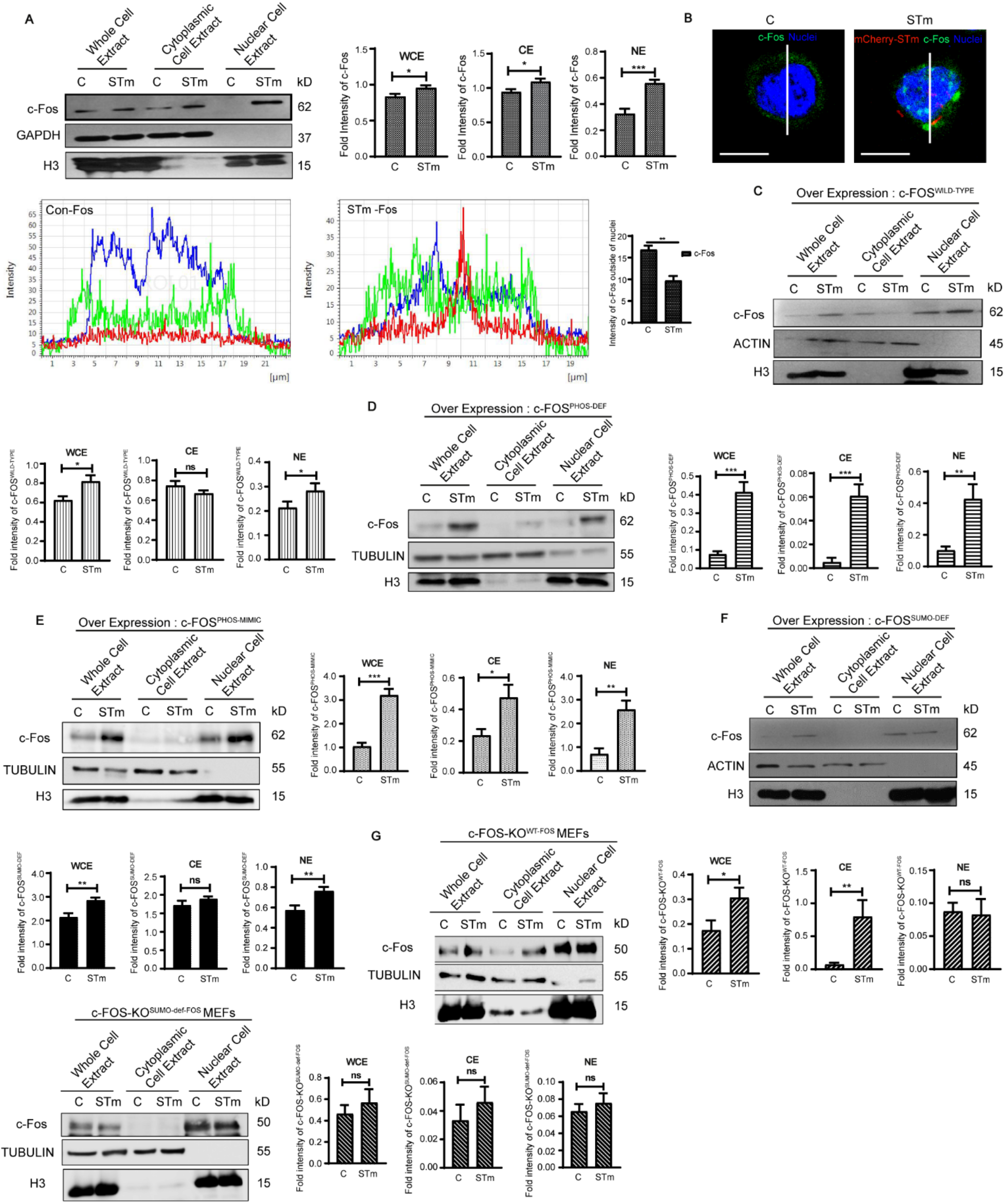
SUMOylation dependent *Salmonella* mediated sub-cellular localization of c-Fos. (A) Fractionation and Immunobloting showing sub-cellular localization of c-Fos in mock infected and 4hrs *STm* infected lysates of HCT-8 cells. (B) z-stack confocal images representing sub-cellular localization of c-Fos (FITC, green) in healthy and 4hrs *STm* (TRITC, red) infected HCT-8 cells. *STm* bears plasmids for mCherry expression and Nucleus was stained with Hoechst (blue). For quantification of cytoplasmic c-Fos expression, in each image, a longitudinal digital line was created across the nuclei of cells (n=10) that defines region of interest (ROI). This was followed by fluorescence intensity measurement of c-Fos within 2µm distance around longitudinal line above of each cells (near drop off intensity of Hoechst at both side). Label distribution graphs showing fluorescence intensity of c-Fos and Nucleus (Hoechst) along cell axis. Distance was shown in pixels (um). (C-F) Fractionation and Immunobloting done for showing sub-cellular localization of c-Fos from HCT-8 cells transfected with plasmid encoding either c-FOS^WILD-TYPE^, c-FOS^PHOS-DEF^, c-FOS^PHOS-MIMIC^ and c-FOS^SUMO-DEF^ in healthy and *STm* infected cells. (G) Immunoblot showing sub-cellular localized expression of c-Fos from healthy and 4hrs *STm* infected lysates of c-FOS-KO^WT-FOS^ and c-FOS-KO^SUMO-def-FOS^ MEFs upon sub-cellular fractionation. For all sub-cellular fractionation experiments specific loading controls were used-GAPDH, Beta-Actin, Tubulin for cytosolic fraction, H3 (Histone3) for nuclear fraction. Densitometric analysis of expression by means + SEM from three independent experiments for the respective protein. Error bars represent SE. Statistical analysis was done using unpaired t-test.

### PTM mediated differential binding of c-Fos to the promoters of target genes

Results presented above suggest that c-Fos mediates a differential regulation of its target genes upon *STm* infection. Various PTMs of c-Fos contribute to this differential regulation. Despite increased nuclear localization of SUMOylation deficient c-Fos (Figure 6F, S7B), we observed a non-uniform change in the gene activation. We thus performed chromatin immuno-precipitation (ChIP) assay to investigate the role of c-Fos and c-Jun PTMs on its capacity to regulate their target genes. HCT-8 cells were either mock infected or infected with *STm* for 4hrs followed by ChIP using c-Fos or c-Jun antibodies. H3 and IgG antibodies were included in the assay as a positive and negative control respectively. Post-immuno-precipitation of the chromatin, quantitative real time PCR based detection (ChIP-qPCR) using oligos corresponding to promoters and non-promoter regions of PIAS1, UBC9 and CCL5 were done. ChIP-qPCR data revealed strong binding of H3 with the immune precipitated DNA which was 18-27-fold above the IgG control in case of all promoters included. The binding of H3 also showed no change in control versus *STm* infected samples. In line with our EMSA data (Figure 2A), a direct binding by c-Fos and c-Jun to the single consensus AP-1 regulatory motifs (TGAGTCA) in the promoters of PIAS1 and UBC9 was observed (Figure 7A, 7B). Binding of c-Fos and c-Jun was also seen in case of CCL5, a known target of AP-1 (Figure 7A, 7B). Upon *STm* infection, c-Fos and c-Jun showed alterations in their abilities to bind to their targets in control and *STm* infected cells. Specifically, binding of c-Fos increased on PIAS1 during infection, while it remained unchanged for UBC9 and CCL5. Strikingly, during *STm* infection compared to control samples, c-Jun showed a decrease in binding to CCL5, UBC9 and PIAS1 (Figure 7B). The differences in the ability of c-Fos and c-Jun to bind to their targets in control and *STm* infected samples led us to examine possible role of PTMs in this mechanism. To examine this possibility, we used custom generated antibodies that specifically bind to SUMOylated form of c-Fos (c-FOS^SUMOYLATED^) described earlier in Tempe et al., 2014 and separately those that bind to phosphorylated c-Fos (FOS^PHOSPHORYLATED^). *STm* infection did not lead to any alteration in binding of c-FOS^SUMOYLATED^ to UBC9 and CCL5, while a small increase in binding in case of PIAS1 was observed (Figure 7C). Interestingly, phosphorylated c-Fos showed strong binding to UBC9 and PIAS1 promoters in the mock infected cells, however, *STm* infection led to a steep decrease ∼3.5 and 1.5-fold respectively. In stark contrast, unlike UBC9 and PIAS1, the binding of phosphorylated c-Fos to promoters of CCL5, CCL19 and CCL2 (highlighted in mRNAseq data) underwent an increase during *STm* infection (Figure 7D). From previous literature, it is known that c-Fos is SUMOylated during active transcription and SUMOylation facilitates its clearance from target promoters (9, 40). In order to understand the exact contribution of PTMs of c-Fos in target gene binding and their transcription, we overexpressed c-Fos in HCT-8 cells, using plasmids that encode a wild type c-Fos (c-FOS^WILD-TYPE^) or a SUMOylation deficient c-Fos (c-FOS^SUMO-DEF^). These cells were either mock infected or *STm* infected for 4hrs followed by ChIP-qPCR (Figure 7E). The data revealed that compared to c-FOS^WILD-TYPE^, overexpression of c-FOS^SUMO-DEF^ showed an increased binding to UBC9 and PIAS1 promoters. Particularly, c-FOS^SUMO-DEF^ showed a dramatic increase in binding (>5 fold) in control versus *STm* infected cells whereas a 2-fold increase was observed in the case of c-FOS^WILD-TYPE^. Thus, these data suggested increased occupancy of SUMOylation deficient variant in SUMO pathway gene promoters and thereby, further highlight the importance of PTMs on c-Fos function.

**Fig 7.**
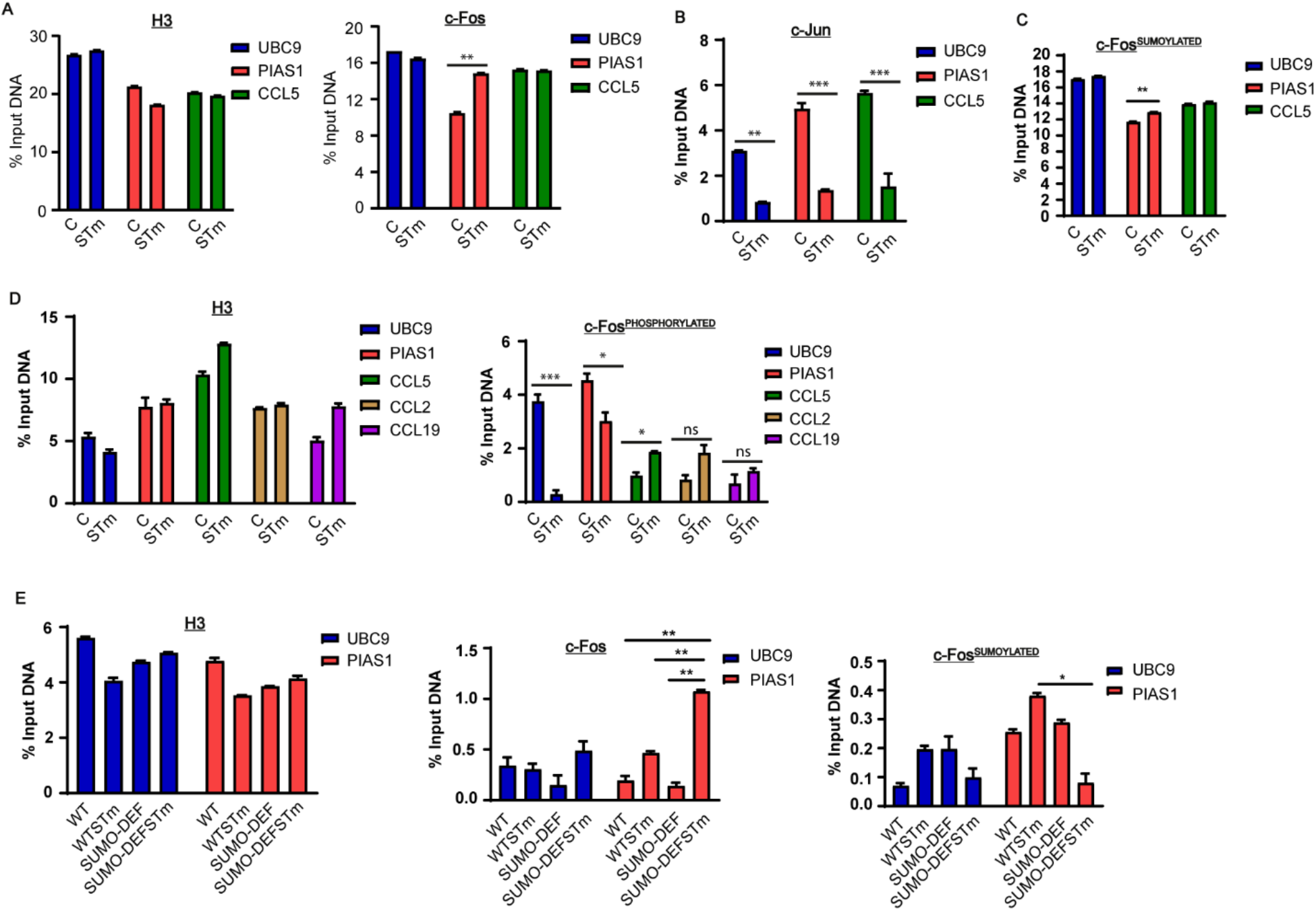
PTMs mediated differential binding of c-Fos to promoters of target genes. (A-B) Chromatin immuno-precipitation assay (ChIP) showing the binding of c-Fos and c-Jun to identified AP-1 regulatory motifs (TGAGTCA) in PIAS1 and UBC9 promoter region at -.893kb and -10.5kb respectively in HCT-8 cells that were either mock infected or *STm* infected for 4hrs. c-Fos, c-Jun, histone3 (H3) and IgG antibodies were used to immune-precipitated the chromatin followed by qRT-PCR for the indicated gene promoters. (C) Antibodies specific for SUMOylated c-Fos (c-Fos^SUMOYLATED^) were used in ChIP (as described in A) to detect binding of SUMOylated c-Fos in mock infected and *STm* infected HCT8 cells. (D) ChIP assay (as depicted in A), using H3 and IgG antibodies and those specific for binding to phosphorylated c-Fos (c-Fos^PHOSPHORYLATED^) to AP-1 regulatory motif in promoter region of UBC9, PIAS1 and CCL2 & CCL19 in mock infected and 4hrs *STm* infected HCT-8 cells. (E) ChIP assay, to detect binding of non SUMOylated and SUMOylated c-Fos to the promoters of UBC9 and PIAS1 genes by using c-Fos^SUMOYLATED^, c-Fos, H3 and IgG antibodies during c-FOS^WILD-TYPE^ and c-FOS^SUMO-DEF^ encoding plasmids in mock infected and 4hrs *STm* infected HCT-8 cells. WT and SUMO-DEF depicts c-FOS^WILD-TYPE^ and c-FOS^SUMO-DEF^ over expression conditions with presence or absence of 4hrs *STm* infection respectively.

**Figure 8.**
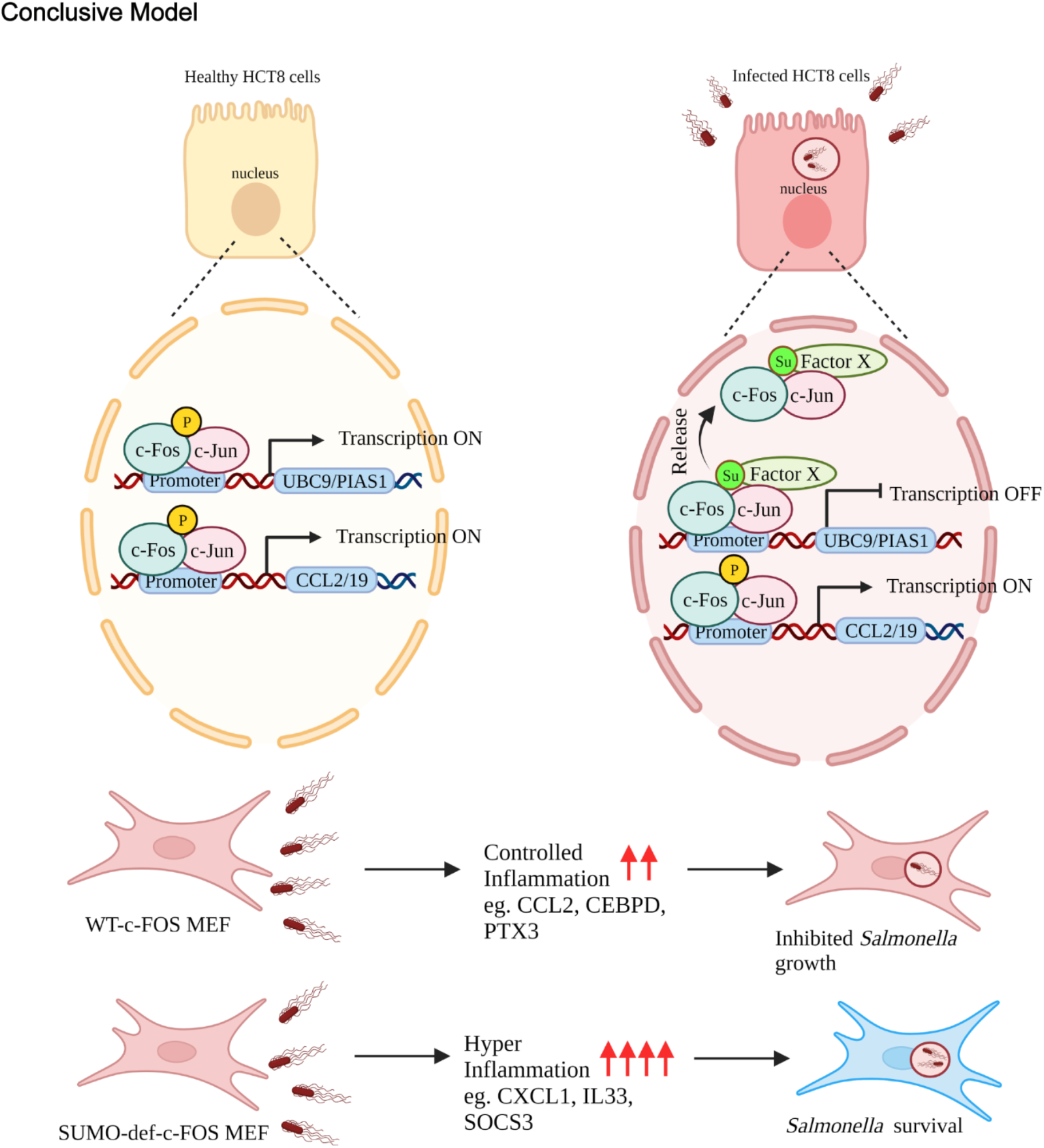
PTM mediated regulation of target genes by c-Fos. Graphical representation of c-Fos function in healthy (top left) and *STm* infected (top right) HCT-8 cells. Image shows turning on of target genes carried out by phosphorylated form of c-Fos due to strong promoter binding. While SUMOylation of c-Fos leads to release of c-For from promoters of SUMO-pathway gens leading to their transcriptional inactivation as seen during *STm* infection. Bottom panel depicts the overall outcome of PTM mediated selective gene regulation, highlighting an immune hyper activation in case of MEFs with a SUMO deficient c-FOS (SUMO-def-c-FOS MEFs) compared to those with wild-type c-FOS (WT-c-FOS MEFs). Such immune dysregulation enables *STm* survival as seen in MEFs with SUMO deficient c-FOS. *Images for this figure have been generated using BioRender (*https://biorender.com/).

## Discussion

SUMOylation machinery is recognised as a major regulator of cellular transcriptional machinery (45). Moreover, altered SUMOylation has been observed in several infections (46) and in inflammatory diseases (47, 48). In an earlier work, we reported a dramatic down regulation of intestinal SUMOylation during *STm* infection (10). Simultaneous down modulation of two key enzymes of SUMOylation pathway, viz Ubc9 and PIAS1 were seen to be required for a successful *STm* infection. SUMOylation of Rab7, a key regulator of intracellular life of *STm*, was shown to be blocked by *STm* (13). Thus, these information point toward the importance of SUMO pathway during bacterial infections. A detailed understanding of the mechanisms of transcriptional regulation of SUMO pathway genes themselves is however lacking. Since, both Ubc9 and PIAS1 undergo down regulation during *STm* infection, the possibility of a common mode of transcriptional regulation was examined in the current study. Our aim was to identify transcription factor(s) that regulate SUMO pathway genes in context of *STm* infection.

Computational tools allowed identification of several putative regulatory motifs in the promoter regions of UBC9 and PIAS1. The prominent motif was that of AP-1 which regulates gene expression of variety of genes in presence of several stimuli including stress, growth factors, cytokines, bacterial and viral infections (29, 49). Early activation of AP-1 is known to occur in macrophages as a result of LPS exposure (50). NF-κB and AP-1 are required for the production of pro-inflammatory cytokine (such as IL-8) during *STm* infection (3). For instance, binding of NF-κB and AP-1 to the CCL2 promoter is known to be responsible for transcriptional activation of CCL2 in pulmonary fibrosis (51). *STm* invasion causes increased expression of CCL2 in HLA-B27 cells (52). Promoter of inflammatory cytokine encoding gene IL-8 contains AP-1 and NF-κB binding protein (CEBP) binding sites. IL-8 gene expression is transcriptionally regulated by AP-1 and NF-κB in presence of *STm* porins and LPS (53, 54). IL-1β treated Caco-2 cells show increased binding of AP-1 on IL-6 promoter responsible for IL-6 production in dose and time dependent manner (55). Here, using EMSA and ChIP assays, we demonstrated a direct binding of c-Fos to PIAS1 promoter. Furthermore, expression of both Ubc9 and PIAS1 was shown to be modulated by AP-1, however a differential regulation is seen when compared to other targets such as CCL2 and CCL19. Presence of potential binding sites of several other factors in the promoters of Ubc9 and PIAS1 identified here may pose interesting areas for future studies.

Reversible PTMs are known to be the main mediators of differential selectivity of targets of transcription factors (26,44,56). A similar example is seen in case of T-bet, a T box transcription factor (T-bet) and a critical modulator for activation of host immune responses. Its activity is mediated by its phosphorylation and ubiquitination. Such a mode of regulation is also seen in case of each of inflammation associated with IL-2, IL-17 and IFNγ (57). GATA3 is known as an important regulator for both adaptive and innate immunity. Phosphorylation, methylation and SUMOylation modifications modulate GATA3 activity in immune cells and regulates IL-4, IL-6 and IL-13 expression. Apart from phosphorylation, O-GlcNAcylation is important for STAT3 mediated IL-10 production and colonic inflammation. AP-1 tightly regulates the expression of cyclin D1 during different phases of cell cycle as a result of its phosphorylation status (58).

Our transcriptome data further highlights differential regulation of several targets of AP-1 by c-FOS and its non SUMOylatable version. This is in particular the case for NOS2, CXCL1, CCL2, FOS, JUN, IRF2, PTGS1, IL-33, IL-6, TNFAIP8 and VEGFA were seen dependent on SUMO-modification of c-Fos. NOS2, a key enzyme of nitric oxide synthesis, was significantly upregulated in *STm* infected c-FOS-KO^SUMO-def-FOS^ in comparison to *STm* infected c-FOS-KO^WT-FOS^. NOS2 is known to play a key role in iron homoeostasis, generation of inflammatory chemokines and cytokines (IL-6 & CCL2) to reduce bacterial numbers in host (59–61). Ligand CXCL11 or its receptor (CXCR3) depletion is also associated with higher bacterial burden in caecum due to compromised intestinal IgA content, CD4^+^ T cells and neutrophils recruitment upon 30-45 days of *STm* infection (62, 63). Upon DSS treatment, upregulated CCL2 expression in CX3CR1 expressing recruited inflammatory macrophages during colitis tissue leads to more severity of inflammatory bowel disease (IBD) (64). CCR2 knockout mice exhibit lower recruitment of iNOS expressing inflammatory monocytes. As a result, nitrate levels are lessened leading to a compromised *STm* infection (65). Similarly, CCL2 another gene identified in our study, plays a critical role for immune response and *STm* survival (66). IRF2 is an essential transcriptional repressor for maturation and development of several immune cells. IRF2 null mice shows impaired NK cells, DCs, Th1 cells and affected B lymphopoiesis and hematopoiesis (67). Cells lacking IRF2 display dampened non canonical inflammasome response upon gram negative bacterial infection (68). LPS or *STm* mediated transcriptional reprogramming causes differential expression of prostaglandins synthesis enzymes (PTGS1) during *STm* infection and control inflammatory response of host via neutrophils chemotaxis (69–71). IL-33 is known as a regulator of gut permeability. IL-33 knock-out mice demonstrates decreased number of paneth cells and fatal systemic *STm* infection. Thus, several of the genes regulated by c-Fos SUMOylation belong to immune pathways, which are known to be specifically involved during infections and intestinal inflammation. It appears that their regulation at the transcriptional level relies on c-Fos and its PTM status. SUMO-modification of c-Fos renders additional level of regulation which is evident from this Quantaseq data, cell fractionation mathematical modeling and ChIP assays. The cell fractionation data reveals that the sub-cellular localization of c-Fos relies on its PTMs, particularly SUMOylation. SUMOylation of c-Fos facilitates clearance of c-Fos from target promoters (9). c-Fos is SUMOylated during active transcription and relives it from target promoters (9). Particularly during *STm* infection, the use of SUMOylated c-Fos specific antibodies in ChIP assays revealed an increased binding at UBC9 and PIAS1 but not CCL5. Decreased binding of phosphorylated c-Fos at UBC9 and PIAS1 promoter was observed upon *STm* infection. This could explain why Ubc9 and PIAS1 show a decreased expression during infection. As represented in the model figure, the selectivity of the regulation of various target genes relies on SUMOylation and Phosphorylation of c-Fos. In a healthy cell, the Phosphorylated c-Fos/c-Jun heterodimer drives the activation of target genes such as UBC9, PIAS1, CCL2 CCL19 and others. On contrary, in a *STm* infected cell, there is a preferential distribution of SUMOylated c-Fos/c-Jun at UBC9 and PIAS1 promoter which turns their transcription off. In infected cells, most of the phosphorylated c-Fos/c-Jun are recruited at CCL2, CCL19 and immune genes to drive their transcription. Such a mode of regulation is an essential step which, if disabled, leads to a compromised intracellular *STm* replication. The current work only focuses on c-Fos and its PTMs but additional layers of complexity might result of PTMs of Jun and other factors and interactors. Studies targeting these would further help our understanding of selective regulation of AP-1 target genes during *STm* infection.

## Materials and Methods

### Materials

All chemicals unless otherwise specified were obtained from Sigma-Aldrich. Antibodies directed against Tubulin (catalog no. T7816), c-Fos (F7799) and Ubc-9 (catalog no. U2634) were obtained from Sigma-Aldrich. Anti-glyceraldehyde-3-phosphate dehydrogenase (anti-GAPDH) (catalog no. 39-8600), anti-c-FOS (catalog no. MA5-15055), anti-MMP7 (catalog no. PA5-28076), anti-c-JUN (catalog no. MA5-15172), Horseradish peroxidase (HRPO)-conjugated anti-rabbit (catalog no. 65-6120) and HRP-conjugated anti-mouse (catalog no. 62-6520) antibodies were obtained from Invitrogen. Anti-H3 (catalog no. ab1791) and anti-c-FOS (phospho T232) (catalog no. ab193365) were obtained from Abcam. Anti-actin (catalog no. 4970), anti-caspase3 (catalog no. 9662), anti-PIAS1 (catalog no. 3550) and anti-c-JUN (phospho S73) (catalog no. 3270) were obtained from Cell Signaling Technology.

### Methods

#### (i) Cell culture

Human colonic adenocarcinoma derived epithelial cell line, (HCT-8) (ATCC, Manassas, VA, USA) (passages 2 to 25) were cultured in RPMI medium supplemented with 14 mMNaHCO3, 15 mM HEPES buffer (pH 7.4), 2 mM glutamine, 1 mM sodium pyruvate, 40 mg/l penicillin, 90 mg/l streptomycin, and 10% fetal bovine serum. c-FOS-KO, c-FOS-KO^WT-FOS^, c-FOS-KO^SUMO-Def-FOS^ f10 MEFs were cultured in Dulbecco’s modified Eagle medium (DMEM) containing 14 mM NaHCO3, 15 mM HEPES buffer (pH 7.5), 40 mg/liter penicillin, 8 mg/liter ampicillin, 90 mg/liter streptomycin, and 10% fetal bovine serum or 0% (serum free) or 20% (switch ON) fetal bovine serum. Initially, c-FOS-KO, c-FOS-KO^WT-FOS^, c-FOS-KO^SUMO-Def-FOS^ f10 MEFs were seeded in 10% FBS containing DMEM media. Post 24hrs cells were starved with serum free media for 12hrs and proceeded for further experiments followed by addition of 20% serum containing DMEM media with or without *STm*.

#### (ii) Bacterial strains, plasmids and infection

*Salmonella* Typhimurium strain SL1344 (obtained from Beth McCormick, University of Massachusetts Medical School, MA) referred to as wild type was used for this study. Strain was grown in Luria broth (LB) at 37°C aerobically for 5 to 7hrs followed by growth under stationary and hypoxic conditions overnight. Overnight cultures were then used to infect HCT-8 epithelial cells at a multiplicity of infection (MOI) of 1:40, while c-FOS-KO, c-FOS-KO^WT-FOS^, c-FOS-KO^SUMO-Def-FOS^ f10 MEFs were infected at a MOI of 1:10. *S*. Typhimurium expressing mCherry was constructed by transforming *S*. Typhimurium with pFPV-mCherry procured from Addgene (Addgene plasmid 20956). Wild-Type c-FOS (c-FOS^WILD-TYPE^) and SUMO-deficient c-FOS (c-FOS^SUMO-DEF^) expression plasmids (27) have been described. Mutagenesis and cloning were carried out to generate phospho-deficient c-FOST232A (c-FOS^PHOS-DEF^) and phospho-mimic of c-FOST232D (c-FOS^PHOS-MIMIC^) using standard PCR-based techniques in c-FOS^WILD-TYPE^ expression plasmid. All details on expression plasmids are available upon request. Flag-JunWT-Myc (Addgene plasmid 47443) were procured from Addgene.

#### (iii) Cell transfection

HCT-8 cells and c-FOS-KO, c-FOS-KO^WT-FOS^, c-FOS-KO^SUMO-Def-FOS^ f10 MEFs were transfected as described earlier (72). One day before transfection, 2.5×10^5^ cells were plated in 24-well plates to obtain 70 to 80% confluency and transfected with Lipofectamine 2000 (Invitrogen, Carlsbad, CA) per the manufacturer’s instructions. Briefly, 0.8 µg of plasmid or 20 pmol of small interfering RNA (siRNA) for c-Fos (Dharmacon ID L-003265-00-0010) or 60 pmol of siRNA for c-Jun (Dharmacon ID L-003268-00-0010) was diluted in Opti-MEM (Invitrogen, Carlsbad, CA). Separately, Lipofectamine 2000 or DharmaFECT (Dharmacon) was also diluted in Opti-MEM and incubated at room temperature for 5 min. Following incubation, the two mixtures were combined and incubated at room temperature for 20 min. This cocktail was added to cells with Opti-MEM and incubated without selection for 24hrs.

#### (iv) In-silico analysis

We carried out an *in-silico* analysis, for each gene, using the Gene Bank Database (NCBI-Gene) a ∼10.6 Kb promoter sequence was extracted. Online software tool, Genomatix: Matinspector was utilized with >0.85 core similarity using EIDorado database 2020 and CiiiDER with 0.15 deficit value using JASPAR database 2020. *In-silico* comparative alignment was done using Gen Bank Database (NCBI-Gene) and Genomatix: Matinspector tool. Multiple sequence alignment study was done using EMBL-MSA: MView Program. Enrichment analysis was done on promoters of SUMO pathway genes with promoters of background genes with p-value p<0.05 using CiiiDER tool.

#### (v) ChIP

Cells were grown in 150mm dish and infected with *S*. Typhimurium. Post infection, 1×10^6^ cells were taken to performed ChIP according to manufacturer kit’s protocol (ab500). The lysed samples were sheared using a sonicator to obtain chromatin fragments of 200–500bp with 60 amplitude 30sec on and 30sec off for 14 cycles. Sheared chromatin was then used for immuno-precipitation using ChIP grade antibodies specific for c-Fos (Invitrogen, MA5-15055), c-Jun (Invitrogen, MA5-15172), SUMOylated c-Fos, phosphorylated c-Fos (abcam. ab193365), H3 (abcam, ab1791) and IgG (abcam, ab171870) isotype acting as negative control overnight at 4°C. The immuno-precipitated DNA was then pulled using DynaBeads^TM^ protein G (Invitrogen) through incubation at 4°C for 3 h on end-to-end rotor. The immuno-precipitated DNA and the input DNA was purified using DNA purifying slurry (Diagenode, C03080030). The purified DNA was used for performing qPCR against Ubc9, Pias1, Ccl2, Ccl19 and Ccl5 promoter. The plates were then run on qPCR machine Bio-Rad CFX 96™. The data were collected, analyzed and plotted using Graph pad prism.

#### (vi) Cell fractionation for EMSA and Immunoblotting

A cell fractionation assay was performed according to protocol (39). Healthy and 4hrs *STm* infected cells were grown in 10 cm dishes. Post-infection, cells were washed with 1xPBS (phosphate buffered saline). Cells were trypsinized and resuspended in 100µl 1xPBS followed by addition of lysis buffer (1xPBS, 0.1% NP40, 1xProtease inhibitor cocktail). Some fraction was aliquoted and labeled as whole cell extract. Remaining lysates were centrifuged for 1 min at 10,000g. Supernatants were collected and labeled as cytoplasmic cells extract. The pellets were resuspended in 1ml of lysis buffer. Supernatant were discarded and the pellet containing nuclear fraction, were solubilized in 0.2 ml of lysis buffer. Equal amount of both cytosolic and membrane fractions were subjected to immunoblotting and EMSA.

#### (vii) EMSA

EMSA was carried out according to DIG gel shift kit, 2^nd^ generation (Roche) kit’s protocol. EMSA was done with small fragment of PIAS1 promoter (39 bp) with nuclear extracted proteins of control cells. The probes corresponding to identify AP-1 binding site on PIAS1 promoter, were 3’ end labeled with Digoxigenin-11-ddUTP (DIG-dd-UTP) molecule by terminal transferase provided in DIG gel shift kit. The probes were incubated with nuclear extract from control cells. For competition assay 100-fold molar excess of unlabeled probes and nuclear extract were incubated along with labeled probes. For Super shift assay, nuclear extract was pre-incubated with anti c-Fos (Invitrogen, MA5-15055) antibody for 2hrs on end-to-end rotor (360°) at 4°C. After pre-incubation all samples were proceeded for binding reaction at room temperature. All samples were run on 5% native polyacrylamide gel.

#### (viii) qRT-PCR

Total RNA was isolated from the samples using the NucleoSpin RNA II kit (Macherey-Nagel, Germany) according to the manufacturer’s protocol. 1µg of each total RNA sample was used to synthesize cDNA using the i-Script cDNA synthesis kit (Bio-Rad, USA). The real-time PCR was performed using a 20µl reaction volume on 96-well plates by using i-Taq Syber Green (Bio-Rad) according to the manufacturer’s instruction in the Bio-Rad CFX 96™ Real Time Detection System. All reactions were normalized to the housekeeping genes GAPDH and HPRT for human and actin and B2M for mouse samples using the cycle threshold (2^-ΔΔ*CT*^) method as described in the detection system. Primer sequences are provided in the supplementary material.

#### (ix) Immunoblotting

*S*. Typhimurium strain SL 1344 was grown and used to infect HCT-8 cells as described above. Following incubation of cells with bacteria for 1 h, the unbound bacteria were washed off and the infection was allowed to proceed for the appropriate time interval. Post infection cells were lysed in Laemmli’s buffer (20 mMTris-HCl pH 8.0, 150 mMKCl, 10% glycerol, 5 mM MgCl2 and 0.1% NP40) separated through a 12% polyacrylamide gel (Bio-Rad, Hercules, CA) by electrophoresis (SDS-PAGE) and transferred to a nitrocellulose membrane. Blots were probed with antibodies against c-Fos, c-Jun, Ubc9, PIAS1, GAPDH, Caspase3, H3, ACTIN, TUBULIN and Mmp7. Blots were visualized by enhanced chemiluminescence using immobilon forte western HRP substrate (Millipore).

#### (x) (a) RNA isolation, 3’mRNA library preparation and sequencing

Total RNA was extracted from 5 million cells using RNeasy Plus Universal Kits (Qiagen, #73404) according to the manufacturer’s protocol. 3’ mRNA specific libraries were amplified using QuantSeq 3’ mRNA-Seq Library Prep Kit FWD (Lexogen, #015.96) using the manufacturer’s instructions with 300ng of extracted RNA. The concentrations of the libraries were estimated using Qubit™ dsDNA HS Assay Kit (Thermo Fisher Scientific, #Q32851). Quality assessment and library size estimation of the individual libraries was done using an HS DNA kit (Agilent, #5067-4626) in a Bio analyzer 2100. The libraries were pooled in equimolar ratio and single-end 100bp reads were sequenced on the Illumina NextSeq 550 platform.

#### (b) Read mapping, counts generation and differential expression analysis

On an average, 7-10 million reads were uniquely mapped per sample. Sequencing quality was assessed using FastQC v0.11.5. Post quality control, the reads were mapped to the *Mus musculus* -mouse *(GRCm38)* reference genome using STAR aligner v.2.5.2a (73). Gene expression levels were measured using the counts generated by HTSeq-count v 0.6.0 (74). To obtain differential expression of the genes, the biological conditions were compared pairwise using DESeq2 (75).

#### (c) Bioinformatics analysis

GO enrichment analysis of the significantly differentially expressed gene set was performed using gProfiler (https://biit.cs.ut.ee/gprofiler/gost) (32). The output of gProfiler was used to generate graphs for KEGG pathways. Read counts were normalized using edgeR (33). Normalized read counts of the genes of interest were fetched from the entire dataset and plotted as heat maps. Heat maps were generated using custom R scripts.

#### (xi) Protein-protein interaction analysis for 3’mRNA sequencing. (a) Data filtration

The proteins with the NaN values are removed from the study. This resulted in the deletion of 35142, 33902, and 36709 genes in the “c-FOS-KO^WT-FOS^ (I) vs c-FOS-KO^SUMO-def-FOS^ (I)”, “c-FOS-KO^WT-FOS^ (UI) vs c-FOS-KO^WT-FOS^ (I)” and “c-FOS-KO^SUMO-def-FOS^ (UI) vs c-FOS-KO^SUMO-def-FOS^ (I)” category respectively.

#### (b) Differently expressed gene (DEG) identification

A fold change cut-off of 1 was taken to identify the DEGs

#### (c) Directional network construction

To create a directional network, we have used the Signor database (76) and extracted the mouse interactome. We only selected those interactions where both the entities are proteins. This resulted in the construction of a directional network (DN) with order 946 and size 1255.

#### (d) Control theory analysis

A dynamical system can be represented as,

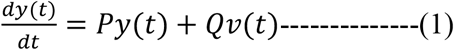

Here, y(t)= [y_1,_y_2_,…………,y_N_]^T^∈ ℝ*^N^* is the state vector of all the N nodes, *P_N_*_×*N*_ is the adjacency matrix of the network, Q=(*b_ij_*) ∈ ℝ*^N^*^×*M*^ (M≤ *N*) is the input vector that contains the nodes that are directly controlled and determines the coupling between the driver nodes and the controlled nodes. v(*t*) = [*v*_1_(*t*), …, *v_M_*(*t*)] ^T^ is the input vector. ℝ denotes the set of real numbers, ℝ*^N^* is the space of real N-vectors, and ℝ*^N^*^×*M*^ is the space of N × M real vectors. To control the entire network, it is important to identify subsets of nodes that, when controlled by distinct signals, are capable of driving the network. These nodes are named as driver nodes in the literature (77–79). According to the controllability rank condition of Kalman (80, 81), the system described by Eq. 1 is controllable if

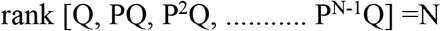

Let G=(V, E), where |V|≠0, be a directed graph, i.e., ∀*e* = (*i*, *j*) ∈ *V*, ∃ a direction from i to j. Here i is called parent node and j is called child node. *M* ⊆ *E* is a matching set if ∄ *e*_1_, *e*_2_ ( *e*_1_ ≠ *e*_2_) ∈ *M* such that *e*_1_and *e*_2_ share a common child or parent node. A matching set of highest cardinality is called a maximum matching. If *e*_2_ = (*i*, *j*) ∈ *M*, then j is called a matching node and all the remaining nodes are called unmatched nodes. In their study, Liu et al. (79) have termed these unmatched nodes as driver nodes and showed that these nodes are sufficient to control the networks. However, the cardinality of many matching sets can be the same, i.e., a network may contain more than one maximal matching set. Hence, the identification of driven nodes is not unique, and several solutions may exist. The network nodes are therefore categorized into critical, intermittent, and redundant categories (82). Critical category contains the nodes which will appear as driver nodes in all the matching sets, intermittent driver nodes are those that appear in some but not all the matching sets while redundant category contains nodes that are not driver nodes. In this study, we are focusing on the identification of critical driver nodes only.

#### (d) Degree centrality analysis

The property of a node being a driver node signifies its global impact. To see the local impact of the nodes in the network, we opt for degree centrality analysis. In a graph G=(V, E), where |V|≠0, the degree of a node *v_i_* is defined as,

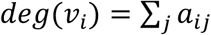

where, A={*a_ij_* } is the adjacency matrix of the network and *a_ij_* = 1 if i and j are connected or 0 otherwise. Hubs are defined as the nodes with the higher degrees in the network. In this study, we have considered a node hub if its degree is twice more than the average degree of the network.

#### (xii) Gentamicin protection assay (GPA)

Cells were infected with *S*. Typhimurium for 1hr in pen/strep antibiotics free 1xHanks balanced salt solution (HBSS) or 20% serum containing DMEM media. The unbound bacteria were then washed off with 1xHanks balanced salt solution (HBSS) followed by incubation in HBSS or 20% serum DMEM media containing 100 µg/ml of gentamicin to kill any extracellular bacteria for 1hr. After 1hr of Gentamicin treatment, cells were incubated with 10 µg/ml of gentamicin until completion of infection. Following incubation, the cells were lysed using 0.1% Triton X-100 in PBS. Samples were serially diluted in sterile PBS and plated onto LB agar plates. CFUs were calculated by counting the colonies obtained the next day. A countable range of 30 to 300 was utilized.

#### (xiii) Immunofluorescence

Cells were grown on cover slips in 24 well plate. For fixed-cell imaging post infection, cells were washed two times in 1xphosphate buffered saline (PBS) and fixed in 3.7% paraformaldehyde pH 7.4 for 15 min at room temperature (RT) followed by washing in PBS. Cells were permeabilized by incubation in PBS containing 0.1% saponin and 1% bovine serum albumin (PSB buffer) for 1hr. For immunostaining, cells were incubated with monoclonal rabbit anti-c-Fos (1:3000, prepared in permeabilized buffer) for overnight at 4° and then washed three times and incubated with Cy5-conjugated goat anti-rabbit IgG (1:400) for a further 2 hrs at RT. Cells were washed three times and incubated in 4,6-diamidino-2-phenylindole (DAPI) (1 µg/ml; Sigma-Aldrich) for 5 min. Coverslips were mounted using prolong diamond antifade (Invitrogen). The cells were observed using a Leica confocal SP8 microscope with a 63×oil objective. Quantification of cytoplasmic and nuclear c-Fos was done using Leica suite X (LAS X) software.

#### (xiv) Statistics

All the results are expressed as the mean ± s.e.m from an individual experiment done in triplicate. Data were analyzed with one-way ANOVA followed by Tukey’s posttest, standard unpaired two-tailed Student’s t test and the Mann–Whitney U-test was used where applicable, with *p* values of 0.05–0.001 considered statistically significant. We evaluated the statistics with GraphPad Prism software.

## Acknowledgments

We thank RCB instrumentation facility. We are thankful for STARS grant (Ministry of Education), ICMR-SRF award of PK, DBT (Department of Biotechnology, Govt. of India) fellowship of PK. We acknowledge Addgene for the plasmids-(20956 and 47443). We also thank Dr Pallavi Kshetrapal for her help.

## SUPPLEMENTARY MATERIALS

**Fig S1.**
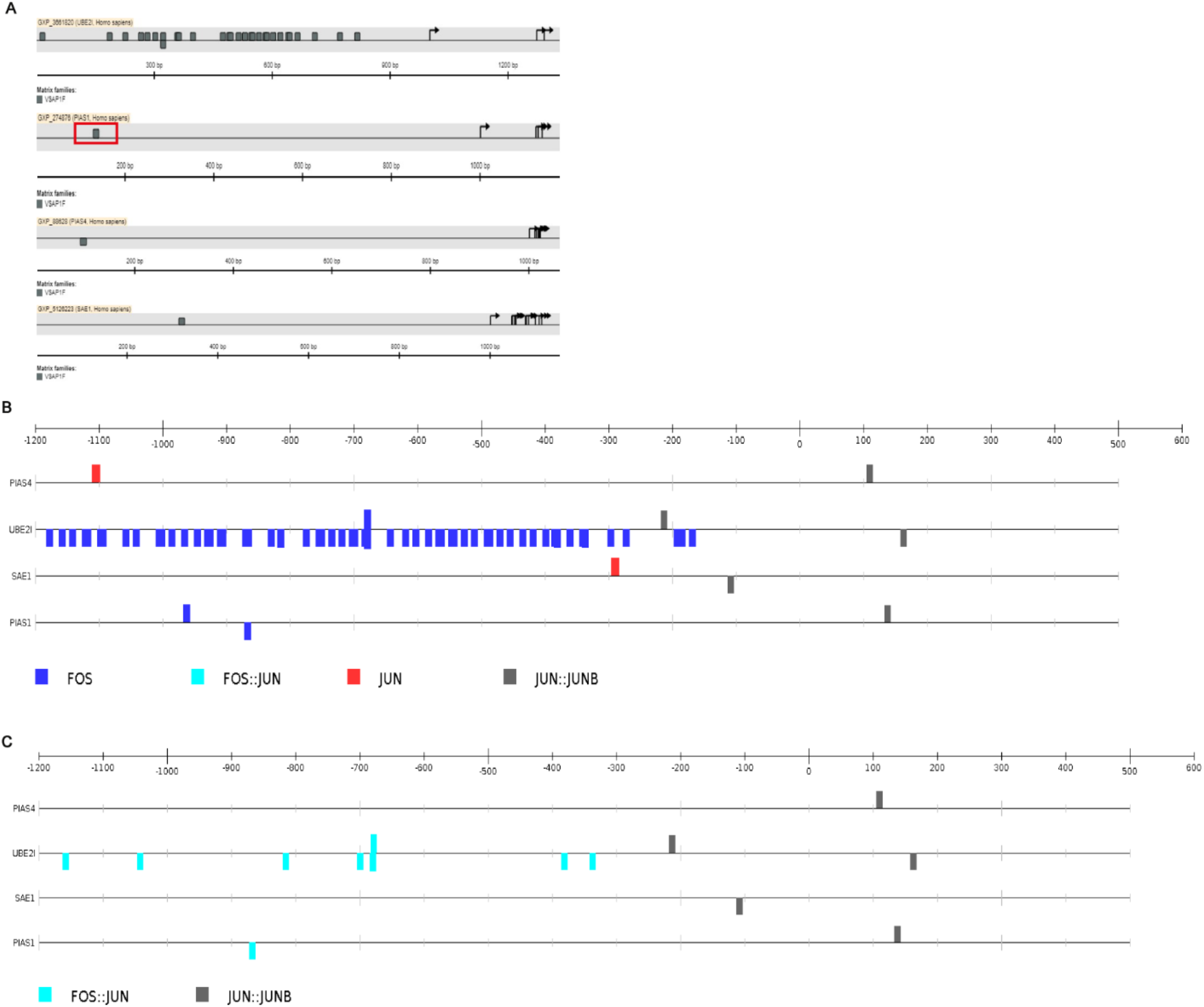
Presence of multiple c-Fos/c-Jun binding motifs in promoters of SUMO-pathway genes. (A) *In-silico* promoter analysis showing existence of consensus AP-1 regulatory motifs (indicated as grey rectangle) in the promoter regions of SUMO pathway genes using Genomatix: Matinspector software tool with >0.85 core similarity using EIDorado 2020 database. (B-C) *In-silico* promoter analysis showing the presence of consensus AP-1 regulatory motifs (along with Homodimer: Jun: Jun, Heterodimer: Fos: Jun) in the promoter of SUMO genes using JASPAR 2020 database based computational CiiiDER tool.

**Fig S2.**
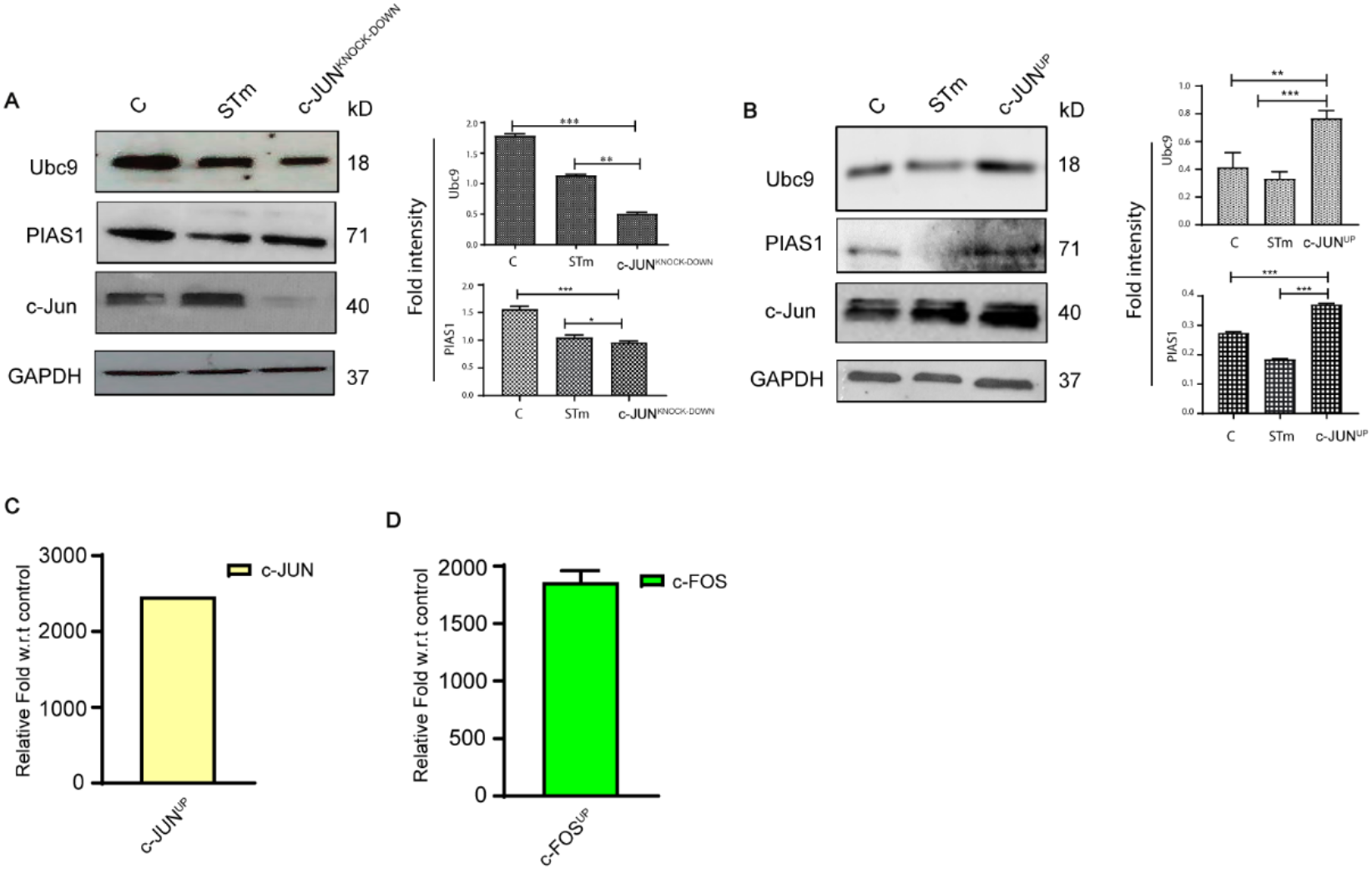
Regulation of expression of Ubc9 and PIAS1 by AP-1 (c-Fos/c-Jun) (A) Immunoblot showing expression of Ubc9 & PIAS1 upon Si-RNA mediated silencing of c-JUN (c-JUN^KNOCK-DOWN^) and 4hrs *STm* infection of HCT-8 cells in comparison to scrambled Si-RNA control. (B) Analysis of expression of Ubc9 & PIAS1 by Immunobloting in HCT-8 cells with c-JUN encoding plasmid based over expression (c-JUN^UP^) and 4hrs infection with *STm* in comparison to untreated control. GAPDH was used as loading control. (C-D) qRT-PCR analysis showing expression of c-Jun and c-Fos upon c-JUN and c-FOS encoding plasmid based over expression (c-JUN^UP^) and (c-FOS^UP^) of HCT-8 cells compare to control sample. Error bars represent mean of SE. Densitometric analysis was done using means + SE of expression data from three independent experiments for the respective protein.

**Fig S3.**
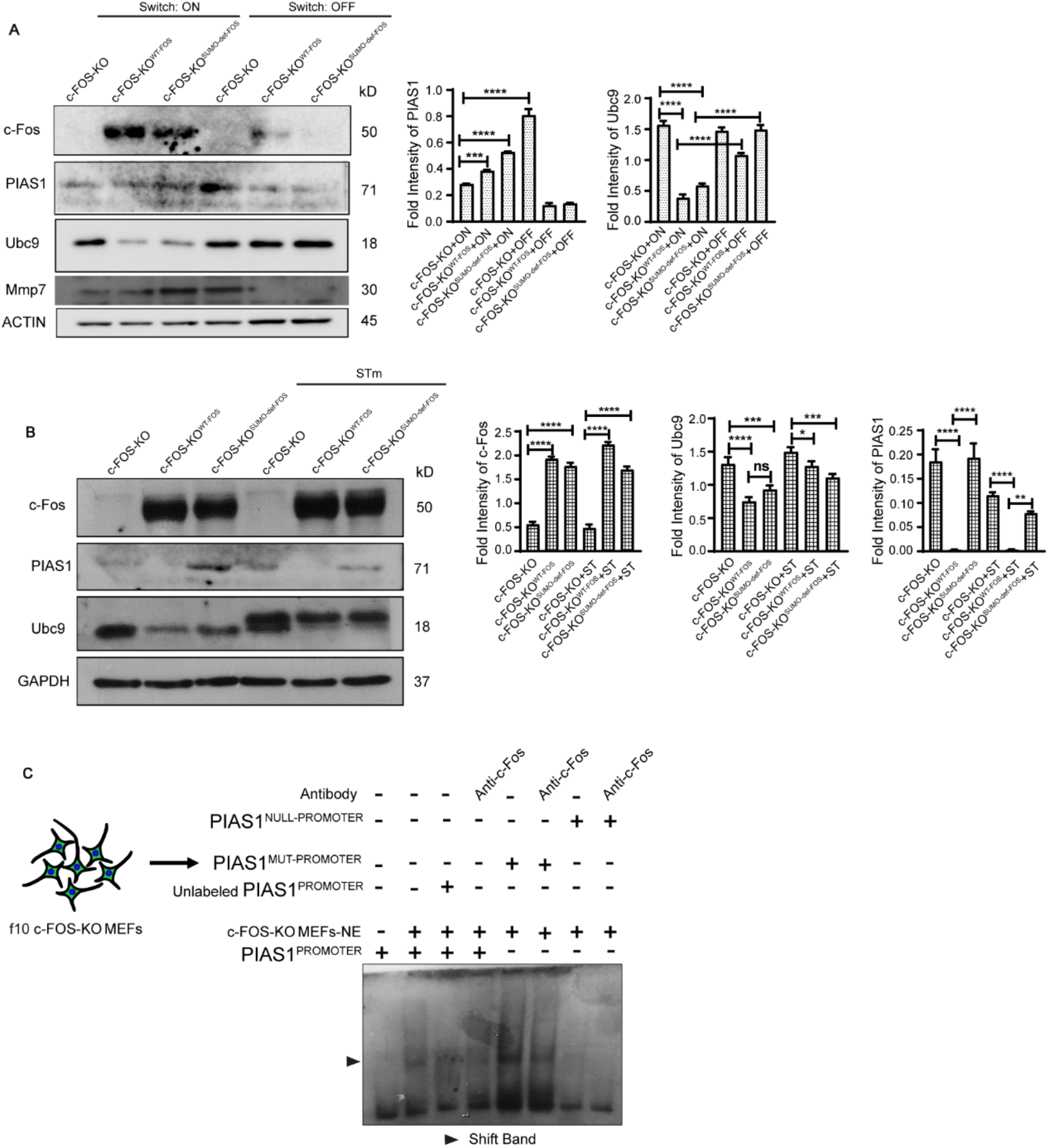
Validation of c-Fos as transcriptional regulator of Ubc9 and PIAS1 genes in c-FOS-KO, c-FOS-KO^WT-FOS^ and c-FOS-KO^SUMO-def-FOS^ MEFs. (A) Immunoblot showing expression of Ubc9 and PIAS1 along with c-Fos during c-Fos induction ‘ON’ and ‘OFF’ conditions in c-FOS-KO, c-FOS-KO^WT-FOS^ and c-FOS-KO^SUMO-def-FOS^ MEFs. Mmp7 was used as positive control gene. (B) Immunoblot showing expression of SUMO (Ubc9 & PIAS1) genes from c-FOS-KO, c-FOS-KO^WT-FOS^ and c-FOS-KO^SUMO-def-FOS^ MEFs in presence and absence of *STm* at 4hrs. (C) EMSA with PIAS1^PROMOTER^, PIAS1^MUT-PROMOTER^ and PIAS1^NULL-PROMOTERs^ was performed using nuclear extract (NE) from healthy c-FOS-KO MEFs. Lane 2, showing presence of shift band (marked with dark arrow) when PIAS1^PROMOTER^ was mixed with nuclear extract (may be due to presence of c-Jun or other protein belonging to AP-1 family). Lane 3, showing absence of shift band in presence of (competitor) 100X unlabeled PIAS1^PROMOTER^, Lane 4, representing absent super shift band in presence of PIAS1^PROMOTER^ and nuclear extract along with anti c-Fos antibody.

**Fig S4.**
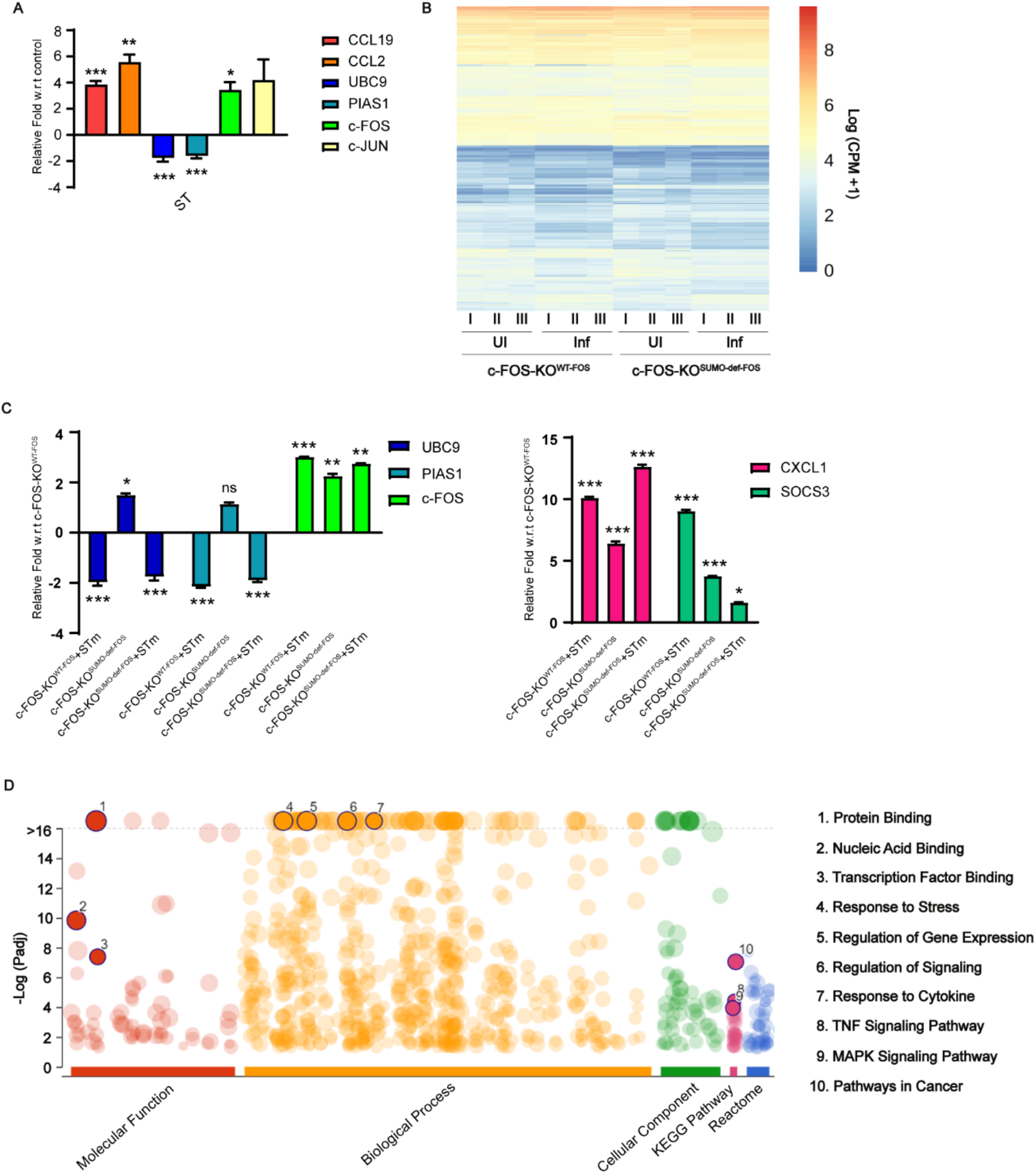
c-Fos SUMOylation mediated immune gene activation and controlled inflammation upon *STm* infection. (A) qRT-PCR analysis showing expression of AP-1, SUMO genes and other AP-1 target genes from HCT-8 cells in presence and absence of *STm* at 4hrs post infection. (B) Heat map showing normalized expression counts of all the significantly differentially expressed genes from different pairwise comparisons from RNA samples of uninfected and 4hrs *STm* infected c-FOS-KO^SUMO-def-FOS^ and c-FOS-KO^WT-FOS^. (C) Validation of shortlisted genes by in-house qRT-PCR. (D) GO analysis of all the significantly differentially expressed genes as generated by gprofiler (https://biit.cs.ut.ee/gprofiler/gost). Few key GO terms highlighted.

**Fig S5.**
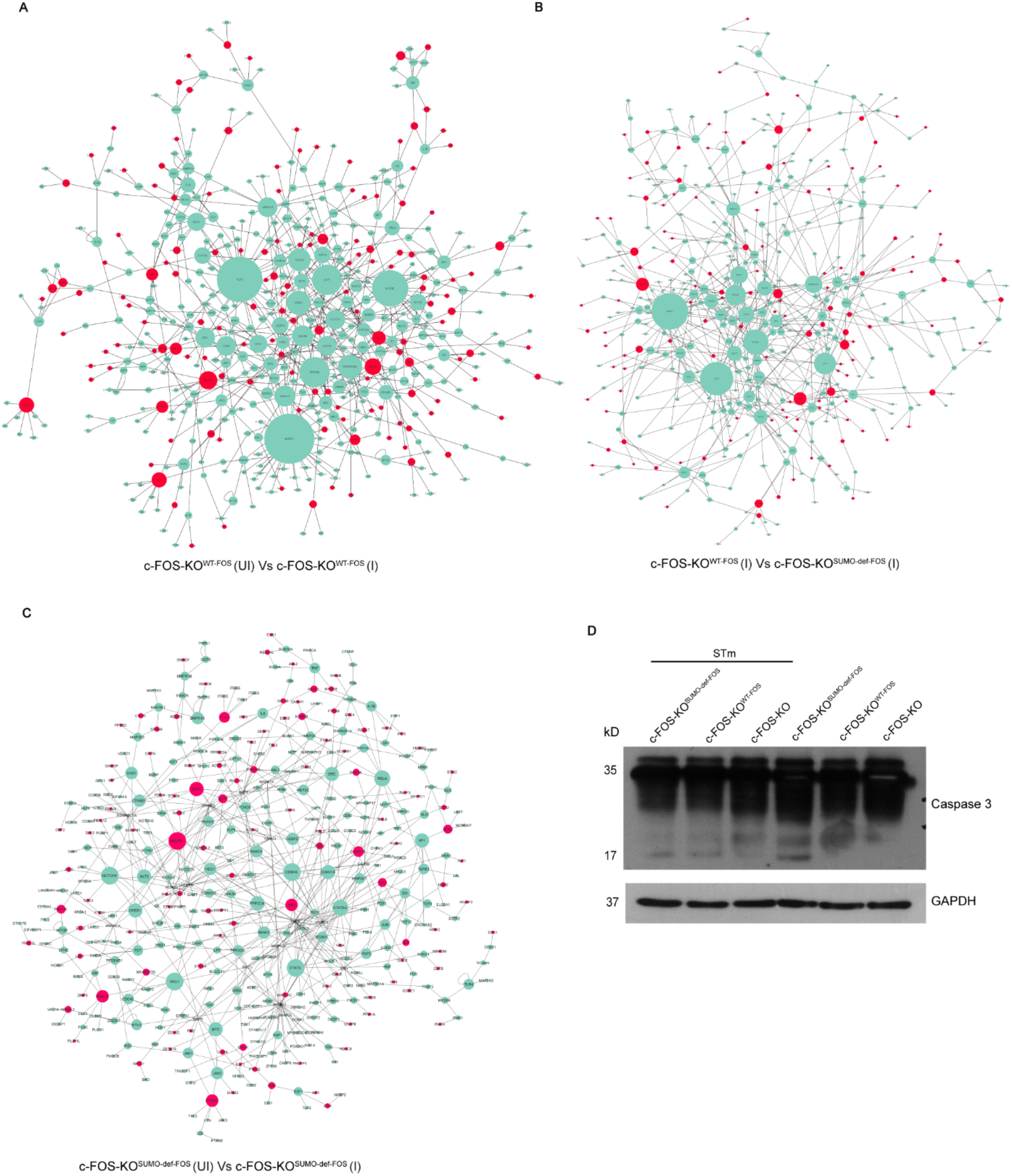
Differentially expressed genes constitute altered protein-protein interaction network and signaling in f10 c-FOS-KO^WT-FOS^ and c-FOS-KO^SUMO-def-FOS^ MEFs upon *STm* infection. (A-C) Constructed mapped proteins (DN) represents as directional network using signor database and extracted mouse interactome from each pairwise comparison c-FOS-KO^WT-FOS^ (UI) Vs c-FOS-KO^WT-FOS^ (I), c-FOS-KO^WT-FOS^ (I) Vs c-FOS-KO^SUMO-def-FOS^ (I) and c-FOS-KO^SUMO-def-FOS^ (UI) Vs c-FOS-KO^SUMO-def-FOS^ (I) during 4hrs of *STm* infection. The network edges are provided in the file network.xlsx. In these figure, red color represents the critical driver nodes and the node size represent the degree. (D) Immunoblot represents activation of caspase 3 in c-FOS-KO, c-FOS-KO^WT-FOS^ and c-FOS-KO^SUMO-def-FOS^ MEFs in absence and presence of 4hrs *STm* infection. GAPDH was used as loading control. Error bars represent SE. Statistical analysis was done using Student’s t-test and one-way ANOVA.

**Fig S6.**
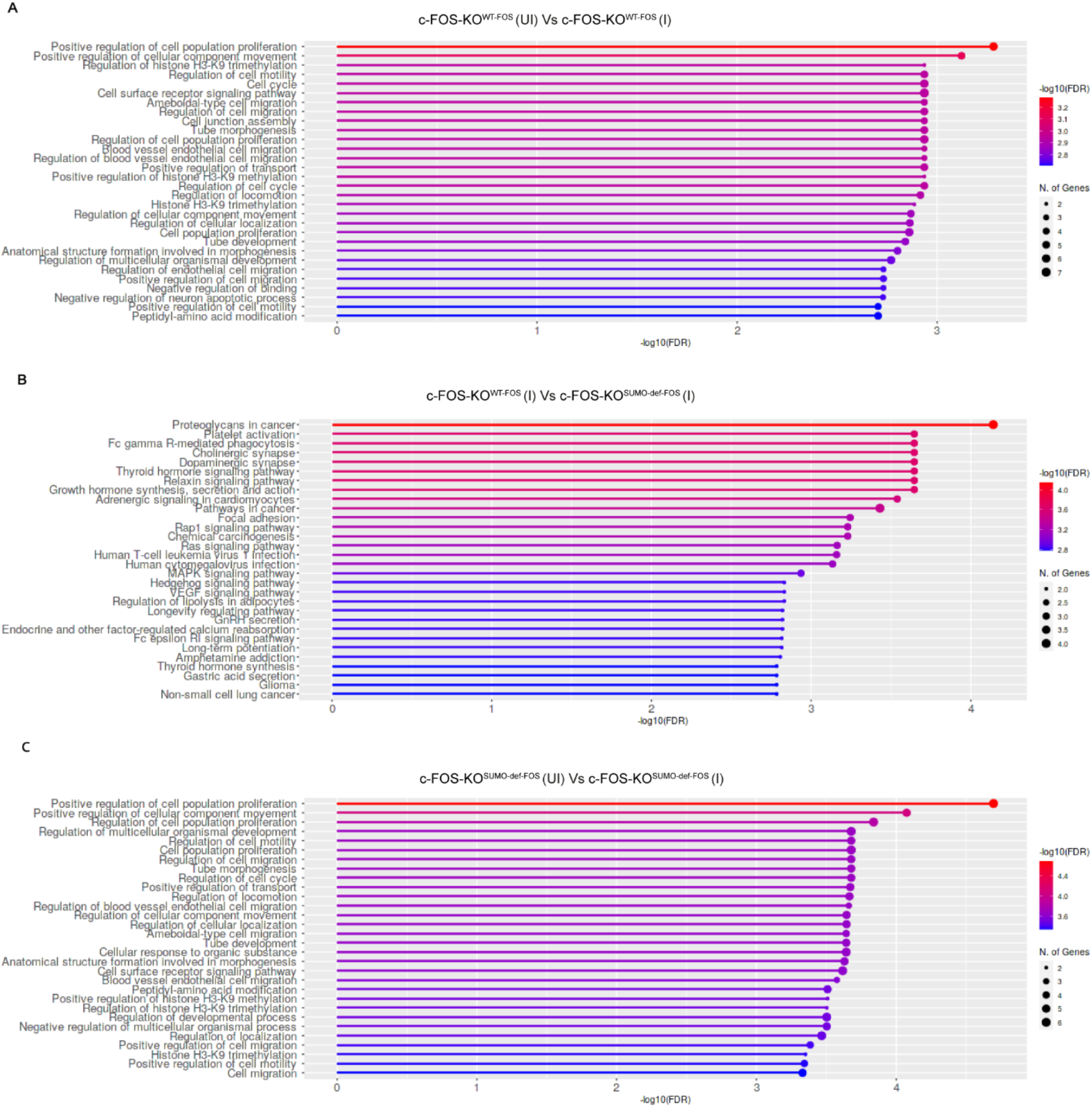
Hub-CDNs are mainly associated with cell proliferation and migration signaling pathway in f10 c-FOS-KO^WT-FOS^ and c-FOS-KO^SUMO-def-FOS^ MEFs upon *STm* infection. (A-C) The crucial KEGG pathways analysis of the hub-CDNs of each pairwise comparison c-FOS-KO^WT-FOS^ (UI) Vs c-FOS-KO^WT-FOS^ (I), c-FOS-KO^WT-FOS^ (I) Vs c-FOS-KO^SUMO-def-FOS^ (I) and c-FOS-KO^SUMO-def-FOS^ (UI) Vs c-FOS-KO^SUMO-def-FOS^ (I) during 4hrs of *STm* infection. Processes are sorted and bars are colored according to the -log10(FDR) values while the size of the dark dots represent number of genes in that process.

**Fig S7.**
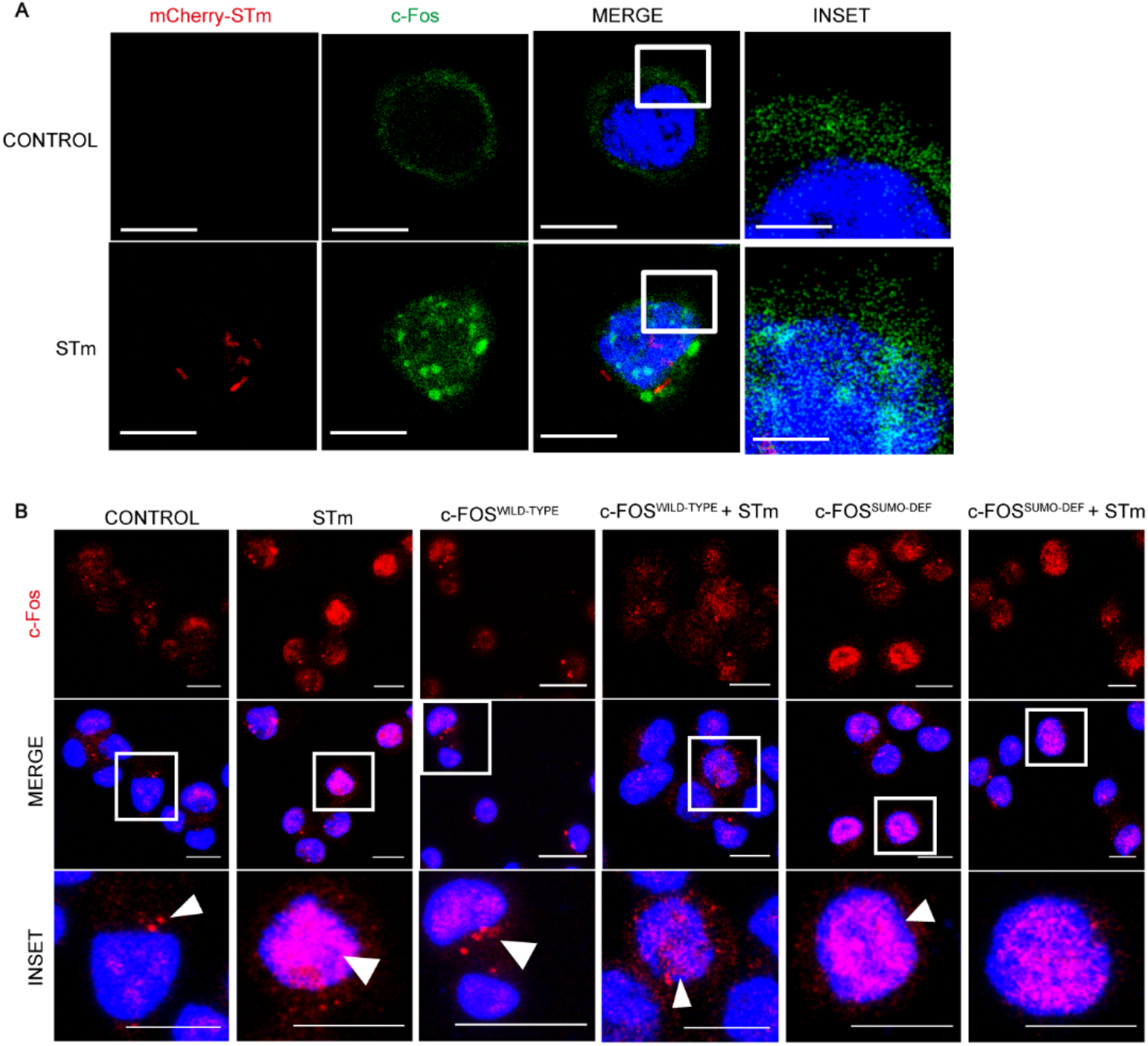
*Salmonella* mediated sub-cellular localization of c-Fos is SUMOylation dependent. (A) z-stack confocal images of sub-cellular localization of c-Fos (FITC, green) in mock infected and *STm* (TRITC, red) infected HCT-8 cells. 4hrs post infection the cells were stained - nucleus (Hoechst, blue) and analyzed. (B) Representative z-stack confocal images of HCT-8 cells transfected with c-FOS^WILD-TYPE^ and c-FOS^SUMO-DEF^ encoding plasmids in mock infected and *STm* infected (4hrs) HCT-8 cells. c-Fos (FITC, red), Nucleus (Hoechst, blue).

### Primers and Promoters list

#### Promoter sequence for EMSA

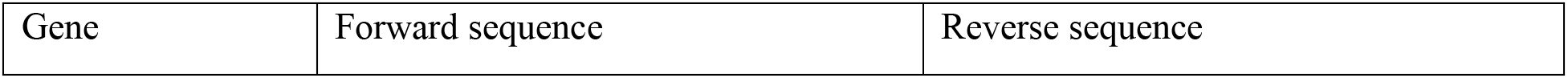
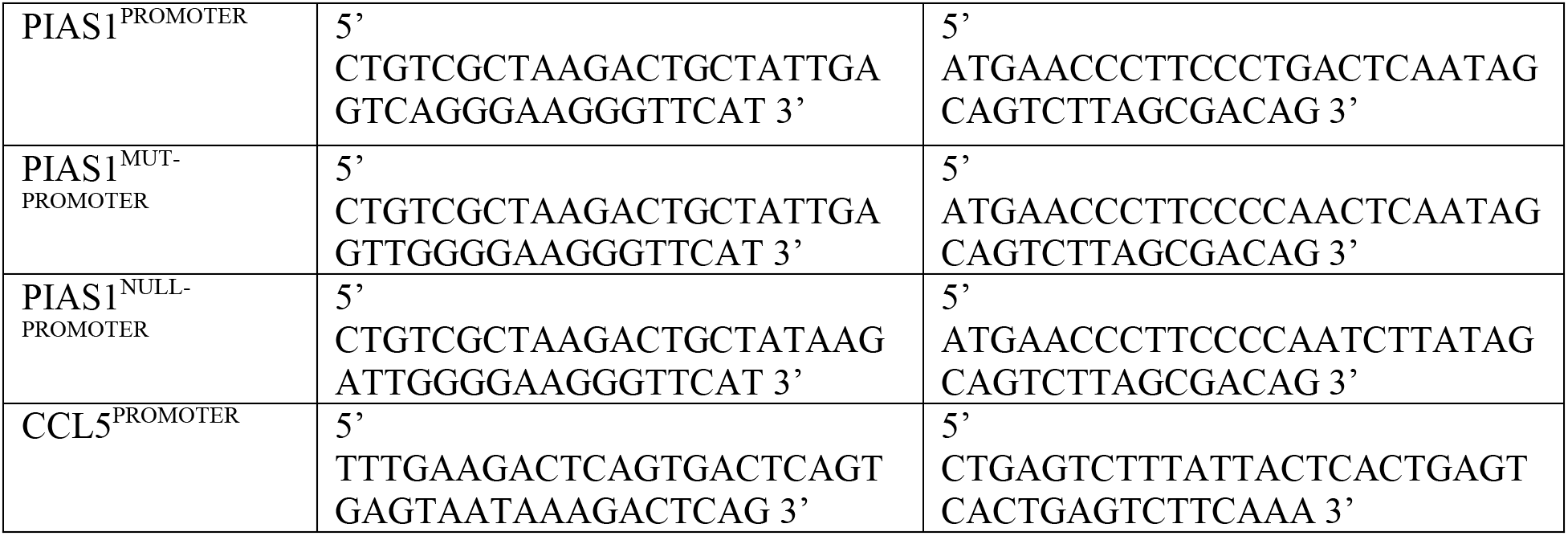

### Primer sequences for RT-PCR

#### Human primers

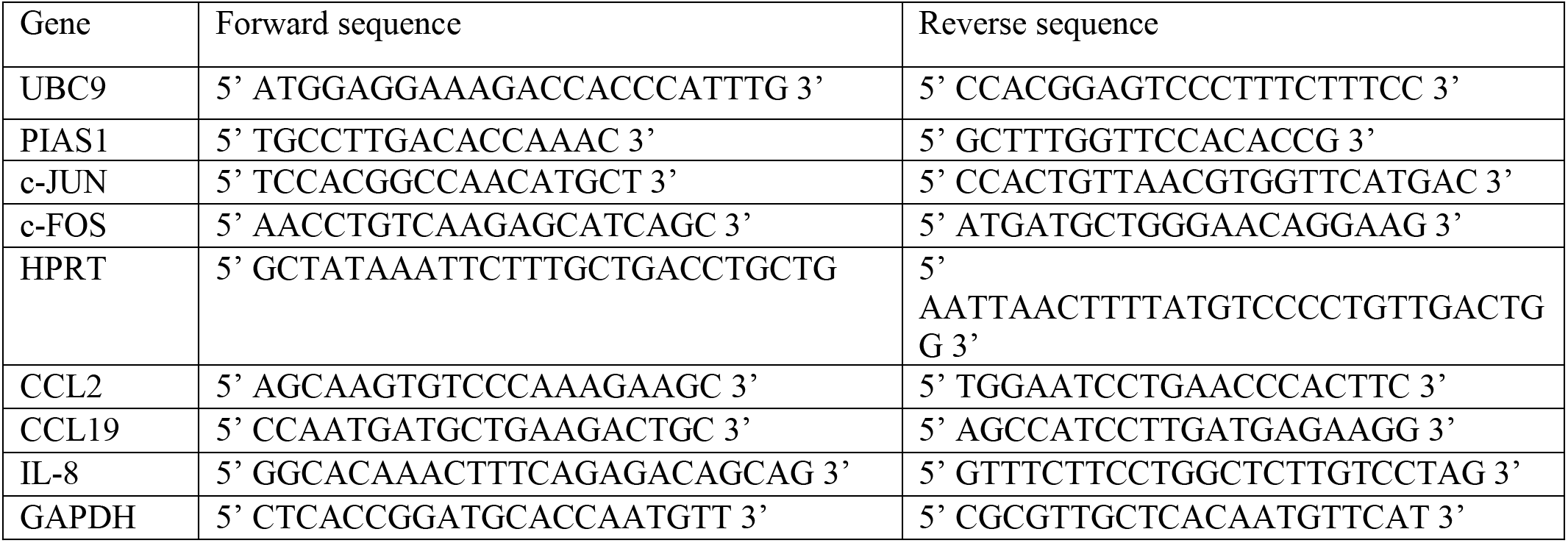

#### Mouse primers

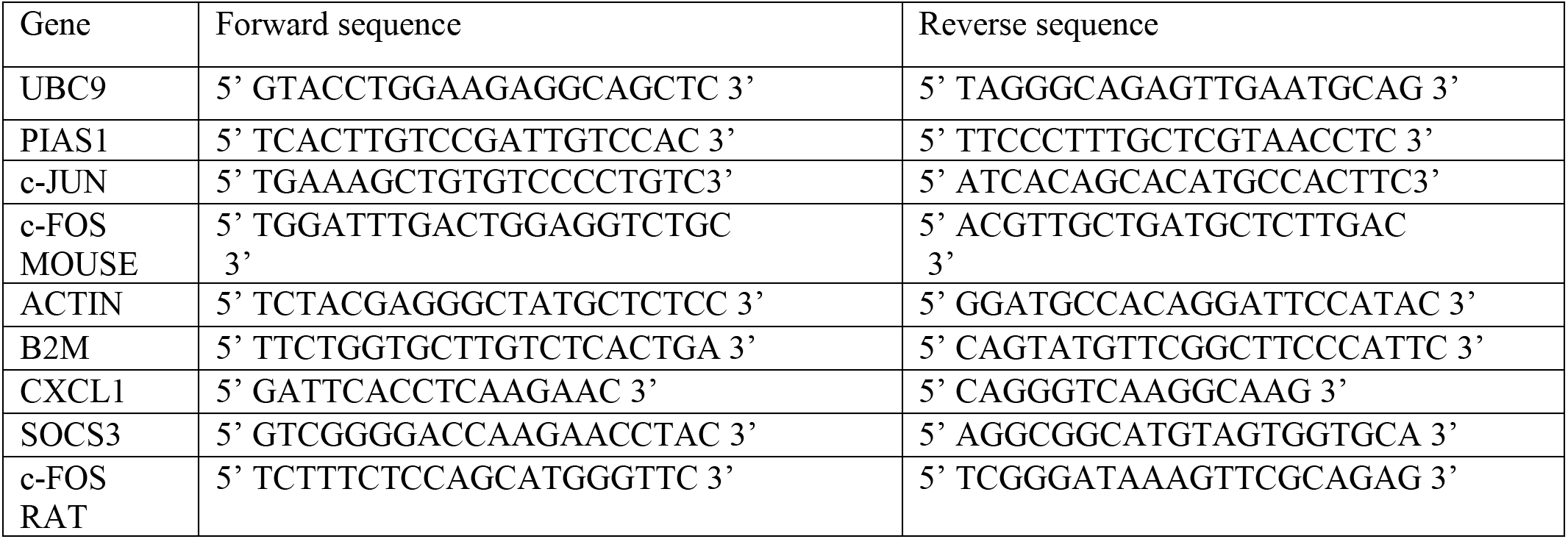

### Promoter sequence for ChIP

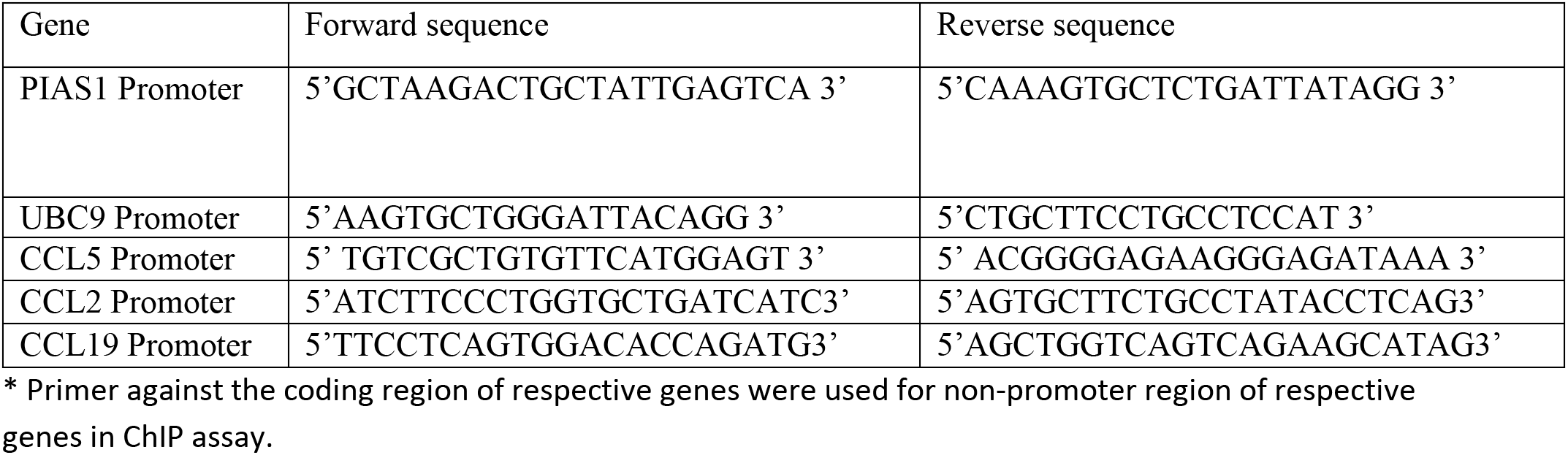

Additional Supplementary information: Additional miscellaneous supplementary files: Gene enrichment analysis, RNAseq-data (XL file), Protein-network analysis data etc

The authors declare no competing interests

